# Alternating hemiplegia of childhood associated mutations in *Atp1a3* reveal diverse neurological alterations in mice

**DOI:** 10.1101/2024.12.24.630241

**Authors:** Markus Terrey, Georgii Krivoshein, Scott I. Adamson, Elena Arystarkhova, Laura Anderson, John Szwec, Shelby McKee, Holly Jones, Sara Perkins, Vijay Selvam, Pierre-Alexandre Piec, Dweet Chhaya, Ari Dehn, Aamir Zuberi, Stephen A. Murray, Natalia S. Morsci, Kathleen J. Sweadner, David A. Knowles, Else A. Tolner, Arn M.J.M. van den Maagdenberg, Cathleen M. Lutz

## Abstract

Pathogenic variants in the neuronal Na^+^/K^+^ ATPase transmembrane ion transporter (*ATP1A3*) cause a spectrum of neurological disorders including alternating hemiplegia of childhood (AHC). The most common *de novo* pathogenic variants in AHC are p.D801N (∼40% of patients) and p.E815K (∼25% of patients), which lead to early mortality by spontaneous death in mice. Nevertheless, knowledge of the development of clinically relevant neurological phenotypes without the obstacle of premature death, is critical for the identification of pathophysiological mechanisms and ultimately, for the testing of therapeutic strategies in disease models. Here, we used hybrid vigor attempting to mitigate the fragility of AHC mice and then, performed behavioral, electrophysiological, biochemical, and molecular testing to comparatively analyze mice that carry either of two most common AHC patient observed variants in the *Atp1a3* gene. Collectively, our data reveal the presence but also the differential impact of the p.D801N and p.E815K variants on disease relevant alterations such as spontaneous and stress-induced paroxysmal episodes, motor function, behavioral and neurophysiological activity, and neuroinflammation. Our alternate AHC mouse models with their phenotypic deficits open novel avenues for the investigation of disease biology and therapeutic testing for ATP1A3 research.

## Introduction

*ATP1A3* encodes the neuronal Na^+^/K^+^ ATPase transmembrane ion transporter necessary to regulate neuronal excitability. Familial and more commonly *de novo* heterozygous variants in *ATP1A3* cause multiple syndromes including alternating hemiplegia of childhood (AHC), rapid-onset dystonia parkinsonism (RDP), and cerebellar ataxia, pes cavus, optic atrophy, and sensorineural hearing loss (CAPOS) with nearly non-overlapping pathogenic variants (Haq et al., 2019; Heinzen et al., 2014; Panagiotakaki et al., 2010; Salles et al., 2021; Tranebjærg et al., 2018). Albeit the heterogeneity in symptoms, patients with pathogenic variants in *ATP1A3* also exhibit common manifestations such as dystonia, seizures, ataxia, and cognitive impairment, indicating the possibility of shared disease mechanisms (Boonsimma et al., 2020; Ishii et al., 2013; Y. Li et al., 2022; Panagiotakaki et al., 2015; Pavone, Pappalardo, Ruggieri, et al., 2022; Rosewich et al., 2012; Salles et al., 2021; Viollet et al., 2015).

Over 1000 individuals with pathogenic variants in *ATP1A3* have been identified of which ∼70% are diagnosed with AHC (Y. Li et al., 2022; Vezyroglou et al., 2022). The most common heterozygous *de novo* missense AHC variants are p.D801N (∼40%), p.E815K (∼25%) and p.G947R (∼10%) that concentrate within or near the transmembrane domains of the ATP1A3 protein (Heinzen et al., 2014; Y. Li et al., 2022; Vezyroglou et al., 2022). AHC patients are affected by sudden and spontaneous episodes (paroxysmal spells), which may be a combination of different symptoms such as hemiplegia, dystonia and seizures (Brashear et al., 2018). Psychological (e.g., excitement, anxiety, anticipation, fright) and environmental stressors (e.g., temperature changes, water exposure) may trigger paroxysmal episodes in AHC patients (Heinzen et al., 2014; Salles et al., 2021). Children and adolescents suffer from the loss of mobility, epilepsies, and complications such as aspiration that can result in early death (Moya-Mendez et al., 2021; Pavone et al., 2022; Salles et al., 2021). Dystonia refers to recurrent episodes of sustained muscle contraction that leads to twisting, repetitive movements or abnormal posture (Van Der Heijden et al., 2022). In general, dystonia is considered a complex neuropathological condition of the motor system involving numerous neuronal circuits of the brain including those of the basal ganglia, thalamus, cortex, brainstem, and cerebellum (Aïssa et al., 2022; Hallett, 2011; Jinnah et al., 2017; Loher & Krauss, 2009; Neychev et al., 2008). Adding to the complexity is the presence of seizures (e.g., focal and generalized seizures), which also underly elaborate neuronal networks (Chauhan et al., 2022; Mueller et al., 2019; Wu et al., 2018). Despite the biological complexity, efforts continue trying to delineate the etiology, involvement of specific brain circuits, and potential long-term consequences that may occur with age and/or because of repetitive paroxysmal episodes in AHC (e.g., neurological regression and deterioration) (Perulli et al., 2022; Saito et al., 2010; Uchitel et al., 2021).

The Na^+^/K^+^ ATPase is a heterotrimeric α-β-FXYD protein complex with one of the four α isoforms (α1 - 4) that are encoded by the *ATP1A1 - 4* genes (Heinzen et al., 2014; Palmgren & Nissen, 2011). The α1 subunit is almost ubiquitously expressed while expression of α2 is restricted (e.g., skeletal, cardiac, and vascular muscle, glia cells, adipose tissues) and α4 subunit expression is observed in male germs cells (Holm et al., 2016; McGrail et al., 1991; Syeda et al., 2020). Expression of the α3 isoform (*ATP1A3*) is observed in excitatory and inhibitory neurons of the mammalian central nervous system including the cortex, hippocampus, striatum, cerebellum, and brainstem (Allen Brain Atlas, Human Protein Atlas, brainrnaseq.org) (Dobretsov et al., 2019; Gokce et al., 2016; Jiao et al., 2022; Smith et al., 2021). In addition, lower ATP1A3 expression has also been observed in the human heart but not in all other species (MGI gene expression data, EMBL single cell expression atlas) (Henriksen et al., 2013; Herrera et al., 1994; McLellan et al., 2020; Premont et al., 2022; Shamraj et al., 1991; K J Sweadner et al., 1994).

ATP1A3 is critical in restoring the transient increase in intracellular Na^+^ concentration after repeated action potentials and in supporting neurotransmitter re-uptake, and thereby, determines the electrical excitability of neurons (Azarias et al., 2013; Dobretsov et al., 2019; Holm et al., 2016; Holm & Lykke-Hartmann, 2016; Zou et al., 2023). Homozygosity for either patient observed variants or loss of ATP1A3 (*Atp1a3* ^-/-^) cause death shortly after birth in mice (Clapcote et al., 2009; Holm et al., 2016; Ikeda et al., 2013). Moreover, heterozygous loss of ATP1A3 (*Atp1a3* ^+/-^) is tolerated in mice without the development of gross morphological, behavioral, dystonia, and survival defects, but causes hyperactivity and enhanced motor locomotion (DeAndrade et al., 2011; Ikeda et al., 2013; Y. B. Liu et al., 2024; Moseley et al., 2007). In stark contrast, mice that are heterozygous for AHC patient associated variants such as p.D801N and p.E815K (referred to as AHC mice) exhibit motor dysfunction, and develop spontaneous and severe stress-induced paroxysmal episodes (Heinzen et al., 2014; Helseth et al., 2018; Hunanyan et al., 2014; Ng et al., 2021; Salles et al., 2021). Interestingly, AHC variants cause reduced ATPase activity, inhibit ATP1A3 pump current, and disturb neuronal excitability. Together with the severe deficits observed in AHC mice, this suggests a potentially dominant-negative disease mechanism for AHC causing variants (Arystarkhova et al., 2019; Heinzen et al., 2012; M. Li et al., 2015; Simmons et al., 2018; Kathleen J. Sweadner et al., 2019).

Several pathogenic AHC variants have been introduced in mice (Clapcote et al., 2009; Hawkins et al., 2024; Helseth et al., 2018; Holm et al., 2016; Hunanyan et al., 2014; Isaksen et al., 2017). The most common p.D801N and p.E815K mutations are associated with early postnatal mortality in mice, and this complicates their experimental investigation. We speculated that hybrid vigor may be sufficient to manage the observed fragility of AHC mouse models. In the presence of hybrid vigor, survival defects were delayed in AHC mice, which allowed us to analyze and directly compare *in vivo* deficits mediated by the D801N and E815K mutations in mice.

## Results

### Hybrid vigor prevents early mortality without abolishing the development of spontaneous and stress-induced paroxysmal spells in B6C3 AHC mice

The two most common dominant *de novo* variants in AHC are p.D801N and p.E815K. The corresponding heterozygous D801N (‘Mashlool’ mouse model; *Atp1a3* ^tm1Ute^) and E815K (‘Matoub’ mouse model; *Atp1a3* ^tm1.1Mika^) animal AHC models have been previously generated and maintained on a congenic C57BL/6J (B6J) background (Helseth et al., 2018; Hunanyan et al., 2014; Uchitel et al., 2021). In addition to the ‘Mashlool’ mouse model, we also introduced the patient-observed D801N variant in C57BL/6J mice (C57BL/6J-*Atp1a3*^em3Lutzy^/Lutzy, Material and Methods, Liu et al., 2024). The B6J D801N and E815K mouse models are highly valuable for ATP1A3 research because mice develop spontaneous and stress-induced paroxysmal spells, which are reminiscent of those observed in patients (Helseth et al., 2018; Hunanyan et al., 2014). Notably, spontaneous death starts after birth and the majority of mutant mice fail to survive past 12 weeks of age (Helseth et al., 2018; Hunanyan et al., 2021; Y. B. Liu et al., 2024). Unfortunately, routine (e.g., breeding, weaning, cage changes) and experimental handling of mice can already induce lethal events (Helseth et al., 2018; Hunanyan et al., 2014), which challenges the ability to maintain a stable and effective ‘live colony’ for either pathogenic variant in B6J mice.

In contrast to pure inbred strains, hybrid vigor allows mice to regain genetic heterozygosity and thereby, are more often resistant to stress and survive better (Birchler et al., 2006; Carol C Linder, 2004; Chan et al., 2017; Selman & Swindell, 2018). Intentionally introducing genetic diversity to mitigate limitations that may arise from inbreeding is a common solution in mouse husbandry, and numerous inbred strains e.g., DBA, C3H, CBA and CAST are viable options for crossing. In the context of seizure traits, C3H (C3H/HeJ) mice are perhaps, particularly interesting because their seizure response greatly differs from that of B6J mice or other inbred strains for various seizure paradigms (Mouse Phenome Database), which led to the discovery of C3H harboring multiple susceptibility and seizure modifier genes (Beyer et al., 2008; Frankel et al., 2014; Tokuda et al., 2009). For example, a benefit of C3H has been observed for other mouse models including Alzheimer mouse models (e.g. APP/PS1 mouse model), which suffer from seizures and mortality when maintained on a congenic B6J background while those defects are absent on a hybrid C3H background (Carlson et al., 1997; Jankowsky et al., 2004; Minkeviciene et al., 2009). Consequently, these strains are available on either genetic background at public strain repositories.

In an attempt to overcome limitations caused by the fragility and early lethality of the B6J AHC mouse models, we crossed the pathogenic B6J D801N and E815K variants onto a C3H hybrid background (referred to as ‘B6C3’, Material and Methods) to generate B6C3.*Atp1a3* ^D801N/+^ (B6C3 D801N) and B6C3.*Atp1a3* ^E815K/+^ (B6C3 E815K) mice. While deaths on a B6J background start after birth with ∼50% of mutant mice failing to even reach wean age (Hunanyan et al., 2021; Y. B. Liu et al., 2024), hybrid B6C3 D801N and E815K mice showed a noticeable onset of unexpected deaths at ∼8 and ∼15 weeks of age, respectively (Figure 1A and B). Mortality of B6C3.*Atp1a3* ^D801N/+^ and B6C3.*Atp1a3* ^E815K/+^ males was generally higher compared to that of mutant female mice (Figure 1A and B). While death of B6C3.*Atp1a3* ^D801N/+^ mice was sudden and spontaneous, only ∼50% of the recorded deaths of B6C3.*Atp1a3* ^E815K/+^ mice were spontaneous (both sexes). The remaining proportion of deaths of B6C3.*Atp1a3* ^E815K/+^ mice required humane euthanasia as a study end point (‘death’) as mice reached an alarmingly low body condition score (BCS) and body weight even more so than B6C3.*Atp1a3* ^D801N/+^ mice (Figure S1A and B).

**Figure 1.**
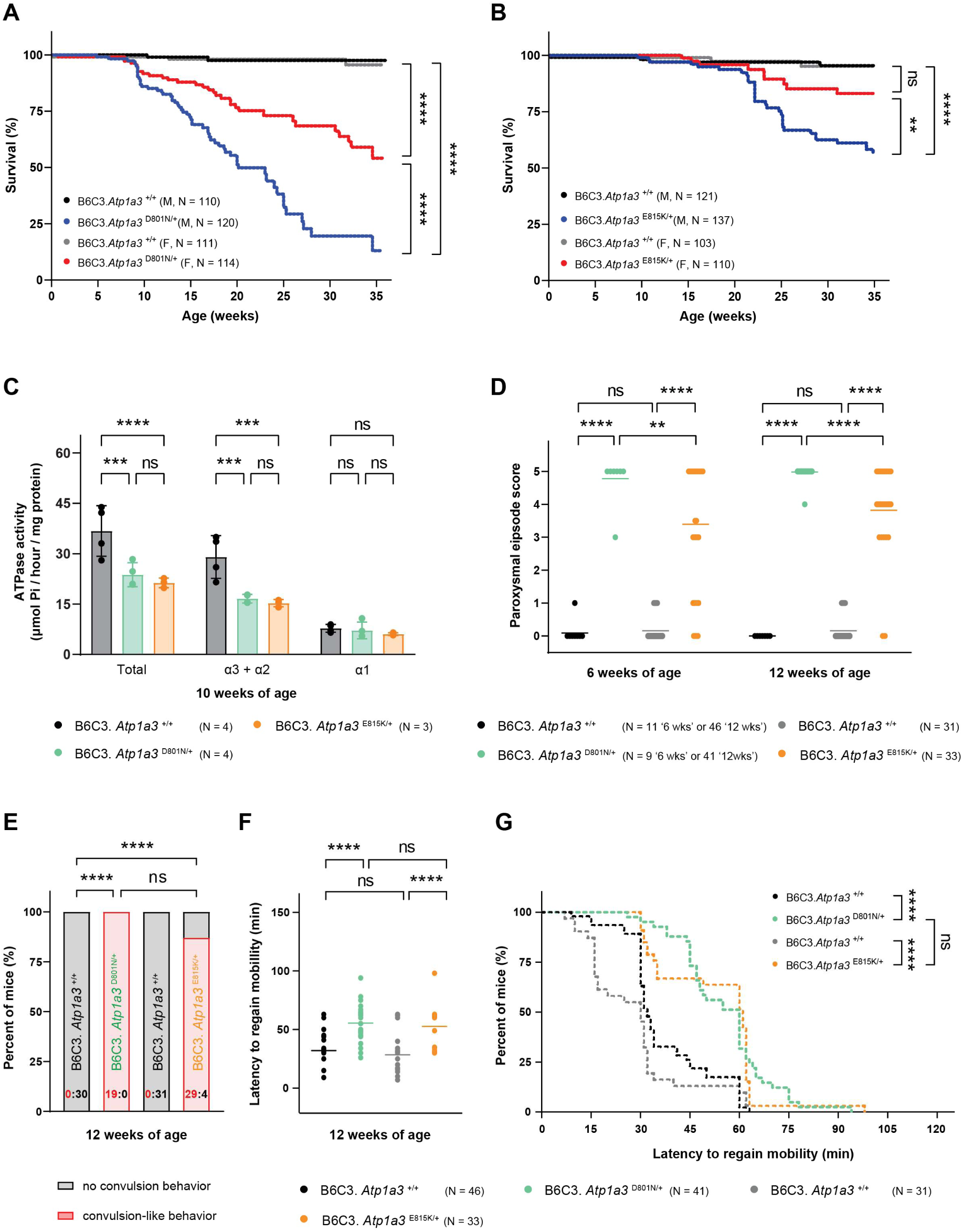
B6C3 AHC mice develop spontaneous and stress-induced paroxysmal spells. (A) Kaplan- Meier survival curve of B6C3.*Atp1a3* ^D801N/+^ mice. Wild type littermate controls for each mutant strain are shown (black and grey). (B) Kaplan-Meier survival curve of B6C3.*Atp1a3* ^E815K/+^ mice. Approximately 50% of B6C3.*Atp1a3* ^E815K/+^ mice required euthanasia as the study end point (‘death’) due to the low body condition score (BCS). Spontaneous and BCS required deaths are included in the survival curve. Wild type littermate controls for each mutant strain are shown (black and grey). (C) Analysis of the ATPase activity of tissues from the hippocampus of B6C3.*Atp1a3* ^D801N/+^ and B6C3.*Atp1a3* ^E815K/+^ mice. Data represent mean ± SD. (D) Hypothermia induced paroxysmal spells (HIP) were induced in B6C3.*Atp1a3* ^D801N/+^ and B6C3.*Atp1a3* ^E815K/+^ mice. Dystonia-like events were scored during the recovery period of HIP experiments and the horizontal line (mean) for each group represents the average score. Wild type littermate controls for each mutant strain are shown. (E) The occurrence of convulsive-like events of B6C3 AHC mice. Data are shown as the fraction (percent) of mice without (grey) or with (red) convulsion-like defects that were observed during the recovery period of HIP experiments. The exact number of mice without (black font) and with (red font) convulsion-like defects is shown in each bar. Wild type littermate controls for each mutant strain are shown. (F) Latency to regain mobility and movement control is shown. The horizontal line for each group represents the mean. Wild type littermate controls for each mutant strain are shown (black and grey). (G) Data in F are shown as a Kaplan-Meier curve. Wild type littermate controls for each mutant strain are shown (black and grey). M, males; F, females; BCS, body condition score. Mantel-Cox test (A, B, G); Two-way ANOVA was corrected for multiple comparisons using Tukey method (C, D); Fisher’s exact test (E); One-way ANOVA was corrected for multiple comparisons using Tukey method (F). ns, not significant; ** *p* ≤ 0.01; *** *p* ≤ 0.001; **** *p* ≤ 0.0001. See also Figure S1.

The ATP1A1 (α1), ATP1A2 (α2) and ATP1A3 (α3) subunits contribute to the total enzymatic Na^+^/K^+^- ATPase activity in the nervous system. We utilized hippocampal tissue samples to assess the ATPase activity in B6C3 AHC mice. The total enzymatic Na^+^/K^+^-ATPase activity was reduced by ∼39% in both B6C3 AHC mice (Figure 1C). The introduction of either the D801N or E815K mutation did not affect the levels of ATP1A3 (α3) expression (Figure S1C and D). In mice, the α1 subunit is much less sensitive to the inhibition by ouabain than α2 and α3, and used to separately assess the activity due to α1 (Y. B. Liu et al., 2024). The α2+α3 ATPase activity was reduced by ∼43% in B6C3.*Atp1a3* ^D801N/+^ and ∼47% in B6C3.*Atp1a3* ^E815K/+^ mice (Figure 1C). The remaining activity of ∼55% in B6C3 AHC mice comprises the ATPase activity that should derive from the unaffected wild type ATP1A3 (α3) subunit and that of the less abundant glial specific α2 subunit. In contrast, the ATPase activity conferred by the ATP1A1 (α1) subunit that is more ouabain-resistant compared to ATP1A3 (α3) and ATP1A2 (α2) was not affected by either AHC mutation (Figure 1C). These data suggest a near complete impairment in the α3 ATPase activity specific to that of the mutant ATP1A3 protein in both B6C3 D801N and E815K mice, which is consistent with previous *in vitro* studies (Heinzen et al., 2012; Weigand et al., 2014).

Patients with AHC suffer from spontaneous spells of symptoms (e.g. dystonia, plegia, and seizures) and B6J AHC mice seem to exhibit similar defects (Helseth et al., 2018; Hunanyan et al., 2014). Handling of mutant mice may be sufficient to induce and observe episodes, which could perhaps, underlie the sudden and spontaneous death in B6J AHC mice (Helseth et al., 2018; Hunanyan et al., 2014). However, we did not witness spontaneous paroxysmal events merely during the routine handling of B6C3 AHC mice. Aiming to better understand the unexpected death of B6C3.*Atp1a3* ^D801N/+^ and B6C3.*Atp1a3* ^E815K/+^ mice, we housed naïve mice in the DIVA (Digital In Vivo Alliance) cage system from ∼13 - 22 weeks of age, which enables continuous video-monitoring of animal activity in their home cage without any operator or experimental interference. We observed that B6C3.*Atp1a3* ^D801N/+^ mice (N = 7) showed a rapid burst in activity including running, jumping and climbing up the wall, just prior to a lethal event that was characterized by freezing, tail and limb twisting, twitching and extension (Supplementary file 1). In contrast, B6C3.*Atp1a3* ^E815K/+^ mice (N = 11) appeared overtly lethargic just prior to their sudden death (Supplementary file 2). Nine of the B6C3.*Atp1a3* ^E815K/+^ mice had an observable episode of freezing and limb twitching (Supplementary file 2) while the remaining mutant mice rather collapsed, which did not allow us to confidently visualize a behavioral episode of freezing and twisting. Although challenged by the fact that the recordings reflect time frames of multiple weeks, we still wondered whether mice would experience multiple spontaneous episodes. Manually inspecting the recordings, we found at least two instances for B6C3.*Atp1a3* ^D801N/+^ (N = 2) and one for B6C3.*Atp1a3* ^E815K/+^ (N = 1) mice in which mutant mice had an episode, appeared to recover and later died because of another episode.

While paroxysmal episodes may not have a clear trigger; stress, exercise, excitement, extreme heat or cold, water exposure, or changes in lighting have been identified to cause episodes in patients with AHC (Heinzen et al., 2014; Salles et al., 2021). Therefore, a variety of conditions to experimentally evoke paroxysmal spells have been previously tested, and hypothermia has been identified as a robust trigger in *Atp1a3* mutant mice (Isaksen et al., 2017; Uchitel et al., 2021). To investigate whether B6C3 AHC mice are also susceptible to this stressor, we adopted the previously described hypothermia paradigm (Figure S1E, Material and Methods) (Isaksen et al., 2017; Pizoli et al., 2002). Mice were placed in 5°C cold water to abruptly induce hypothermia with a body temperature reduction of at least 10°C (Figure S1F). As expected (Hunanyan et al., 2021; Isaksen et al., 2017), no paroxysmal episodes were observed in wild type mice (Figure 1D). In contrast, hypothermic B6C3.*Atp1a3* ^D801N/+^ mice consistently developed paroxysmal episodes with severe dystonia-like postures, lack of voluntary movement, and hyperextended, cramped and stiffened extremities (e.g., legs, paws and/or tail) during the recovery period (Figure 1D, Figure S1G). In agreement with previous hypothermia studies (Hunanyan et al., 2021; Isaksen et al., 2017), spontaneous bursts of ‘convulsion-like’ movements were also observed in B6C3.*Atp1a3* ^D801N/+^ mice (Figure 1E, Supplementary file 3), suggesting that mice may experience a combination of symptoms similar to AHC patients. B6C3.*Atp1a3* ^E815K/+^ mice also developed paroxysmal episodes with dystonia-like postures but had a significantly greater range in severity with some of the B6C3.*Atp1a3* ^E815K/+^ mice even failing to show any abnormal behavior (Figure 1D, Figure S1G). In addition, no ‘convulsion-like’ events were observed in these ‘non-responding’ B6C3.*Atp1a3* ^E815K/+^ mice (Figure 1E). Previous studies noted that the penetrance of paroxysmal spells may be ∼50% or more in hypothermic B6J D801N (dystonia- and convulsion-like events) and ∼65% in forced swim tested B6J E815K mice (dystonia- but no convulsion-like events) (Helseth et al., 2018; Hunanyan et al., 2021). Importantly, hypothermic B6C3.*Atp1a3* ^D801N/+^ (complete penetrance) and hypothermic B6C3.*Atp1a3* ^E815K/+^ (∼90% penetrance) mice developed paroxysmal episodes characterized by dystonia- and convulsion-like body postures, highlighting that these clinically relevant deficits are robust in our hybrid vigor B6C3 AHC models.

Furthermore, paroxysmal spells severely impaired voluntary movement and mobility of B6C3 AHC mice and concordantly, mutant mice had a significant delay until regaining movement control and returning to normal behavior compared to wild type mice (Figure 1F and G). In addition to the data shown (Figure 1F and G), we initially worked with B6C3.*Atp1a3* ^D801N/+^ mice (> 50 mice) to establish and validate the hypothermia protocol. Collectively, less than 3% of AHC mice failed to recover during testing and instead died because of apparent lethal hypothermia-induced paroxysmal spells. B6C3 AHC mice also showed a delay in body temperature recovery (Figure S1F). Although we cannot rule out a link between paroxysmal spells and thermoregulation in mutant mice, we speculate that the delay in body temperature recovery may rather be influenced by the lower body weight of the mutant mice (Figure S1A and B).

### B6C3 AHC mice exhibit impaired motor function and altered behavioral activity

Adding to the dysfunction of the motor system that is evident by the development of dystonia, AHC patients also suffer from ataxia and that in turn, impedes their ability to balance, stand, walk, and run. During routine handling, we noticed that B6C3.*Atp1a3* ^E815K/+^ mice developed an unsteady gait, and it became visually notable at ∼12 weeks of age (Supplementary file 4). Therefore, we subjected B6C3 AHC mice to rotarod testing to assess their motor function. B6C3.*Atp1a3* ^D801N/+^ and B6C3.*Atp1a3* ^E815K/+^ mice showed reduced rotarod performance compared to that of wild type mice, further supporting the impairment of motor function in both B6C3 AHC models (Figure 2A). Interestingly, B6C3.*Atp1a3* ^D801N/+^ mice retained a better rotarod performance with age compared to B6C3.*Atp1a3* ^E815K/+^ mice (Figure 2A). Motor performance not only reflects baseline motor activity but also motor learning (‘modification of motor activity’), which occurs as a result of ‘practicing’ and internal changes in the motor networks (Baladron et al., 2023; Cording & Bateup, 2023; Kogan et al., 2023; Seidler, 2010). Gross motor activity (baseline) and motor learning can be simultaneously or selectively affected (Cording & Bateup, 2023; Duchon et al., 2011; Levin et al., 2006; Pass et al., 2022; Verslegers et al., 2015; Vo et al., 2018; Wagner et al., 2019). While wild type mice improved their rotarod performance over consecutive testing days as expected, neither B6C3.*Atp1a3* ^D801N/+^ nor B6C3.*Atp1a3* ^E815K/+^ mice improved their performance, indicating that motor learning is also impaired in these mice (Figure S2A).

**Figure 2.**
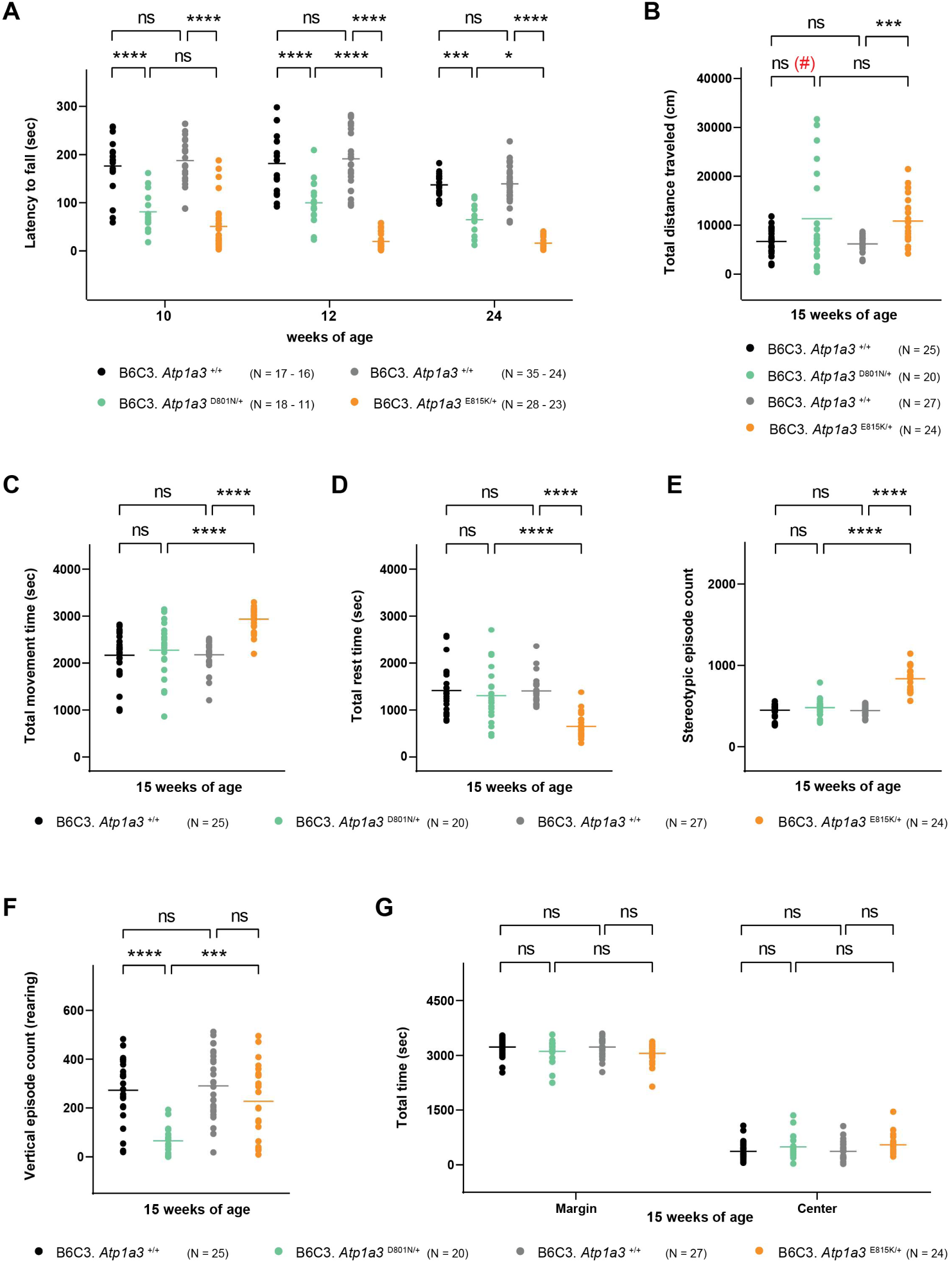
B6C3 AHC mice show motor and behavioral deficits. (A) Latency of B6C3 AHC mice to fall off the accelerating rotarod at various ages. The horizontal line for each group represents the mean. Wild type littermate controls for each mutant strain are shown (black and grey). (B) to (G) Naïve B6C3 AHC mice were subjected to open field testing to interrogate exploratory and/or locomotor activity. The horizontal line for each group represents the mean. Wild type littermate controls for each mutant strain are shown (black and grey). (B) Total distance traveled. Note: # D801N mice only showed a “significant” difference in total distance traveled when data were not corrected for multiple comparisons. *p* = value of 0.0051 compared to littermate control mice (black) or *p* = value of 0.0018 when compared to wild type mice (grey). (C) Total movement time. (D) Total rest time. (E) Stereotypic episodes. (F) Vertical episodes. (G) Total time spent in the center or margin of the open field arena. Two-way ANOVA was corrected for multiple comparisons using Tukey method (A, G); One-way ANOVA was corrected for multiple comparisons using Tukey method (B, C, D, E, F). ns, not significant; * p ≤ 0.05; *** p ≤ 0.001; **** p ≤ 0.0001. See also Figure S2.

Intrigued by the improved survival, motor system deficits, and the ability to handle B6C3 AHC mice and because mild-to-moderate social and cognitive impairments have been observed in patients with AHC (Pavone et al., 2022; Polanowska et al., 2018; Uchitel et al., 2020), we investigated whether additional behavioral alterations can be detected in B6C3 AHC mice. We assessed the grip strength of B6C3.*Atp1a3* ^D801N/+^ and B6C3.*Atp1a3* ^E815K/+^ mice. B6C3 AHC mouse models exhibited a reduction in grip strength (Figure S2B). However, no grip defects were observed when normalized to the body weight of mice (Figure S2B).

Moreover, naïve B6C3.*Atp1a3* ^D801N/+^ and B6C3.*Atp1a3* ^E815K/+^ mice were subjected to open field testing, which assesses changes as a function of exploratory and/or locomotor activity since mice are allowed to freely navigate within a novel environment (Carter & Shieh, 2015; Seibenhener & Wooten, 2015; Tatem et al., 2014). B6C3.*Atp1a3* ^E815K/+^ mice showed a significant increase in the total distance traveled while B6C3.*Atp1a3* ^D801N/+^ mice showed a notably greater range in the distance traveled, which was only significantly increased when not correcting for multiple comparison (Figure 2B). Furthermore, B6C3.*Atp1a3* ^E815K/+^ mice also showed an increase in the movement time and correspondingly, a decrease in the rest time (Figure 2C and D). In addition, the number of stereotypic episodes, which typically reflect repetitive behavioral patterns (e.g., head bobbing, grooming) was also increased in B6C3.*Atp1a3* ^E815K/+^ mice (Figure 2E). In contrast, the movement time, rest time and stereotypic episodes of B6C3.*Atp1a3* ^D801N/+^ were comparable to wild type mice (Figure 2C, D and E). While B6C3.*Atp1a3* ^E815K/+^ mice showed changes in horizontal activity (e.g., movement and rest time), we observed a reduction in vertical activity (rearing) of B6C3.*Atp1a3* ^D801N/+^ mice (Figure 2F). However, neither B6C3.*Atp1a3* ^D801N/+^ nor B6C3.*Atp1a3* ^E815K/+^ mice exhibited significant differences in the time spent in the center or the outer zone (margin) of the open filed arena in which changes could be indicative of anxiety-related behaviors (Figure 2G). Although activity changes in the open field arena can be impacted and correlated with physical and/or locomotor deficits (e.g., ataxia, tremor, muscle weakness); the directionality of the observed activity changes in D801N (reduction in vertical activity) and E815K (increase in horizontal activity) mice failed to tie in with the motor, body weight and grip strength deficits (Figure 2A, Figure S1A and B, and Figure S2B). Alternately, the differential changes in open field activity may instead reflect functional changes in exploratory (e.g., cognitive) behavior of B6C3 AHC mice.

Some patients with AHC and *ATP1A3*-related syndromes show bradycardia and cardiac rhythm abnormalities (e.g., shortened QTc intervals) (Balestrini et al., 2020; Jaffer et al., 2015; Moya-Mendez et al., 2021). Pathogenic variants in *ATP1A3* may directly or indirectly impair heart function through the excessive excitability of the brain (Aiba & Noebels, 2015; Balestrini et al., 2020; Hunanyan et al., 2014). However, no significant electrocardiogram changes (e.g., interval and amplitude) were observed in B6C3 AHC mice (Figure S2C).

### B6C3 AHC mice exhibit altered hippocampal and cortical network activity

Patients with AHC, especially those with the pathogenic p.E815K variant may, develop severe seizures and epilepsy (Capuano et al., 2020; Ford et al., 2023; Panagiotakaki et al., 2015; Viollet et al., 2015). Previous electrophysiological recordings in hippocampal slices of B6J AHC mice revealed an increase in neuronal excitability and may contribute to the observed seizure susceptibility in patients (Clapcote et al., 2009; Hunanyan et al., 2018). Using cortical (visual cortex, V1 and motor cortex, M1) and hippocampal (dorsal CA1 area) alternating current local field potential (AC-LFP) and direct current (DC)-potential recordings in freely behaving wild type, B6C3. *Atp1a3* ^D801N/+^ and B6C3.*Atp1a3* ^E815K/+^ mice (Figure 3A), we searched for possible epileptiform activities and other alterations in neuronal activity in the brain of B6C3 AHC mice at various ages (e.g., AC-LFP trace examples at 14 weeks of age, Figure 3B). Consistent with the influence of the C3H genetic background (Frankel et al., 2014), spike-wave discharges (SWD) were observed in the M1 cortex of B6C3 wild type mice but were much less frequent in B6C3 AHC mice (Figure 3C). The rate of hippocampal spikes increased in B6C3.*Atp1a3* ^E815K/+^ mice with age up to ∼65 spikes/hour at 14 weeks of age (Figure 3D) while in B6C3.*Atp1a3* ^D801N/+^ the rate was similar to that in wild type mice and remained constantly low over the different ages (Figure 3D). Furthermore, only B6C3.*Atp1a3* ^E815K/+^ mice showed hippocampo-cortical giant spikes that are epileptiform features predictive of seizures (Gureviciene et al., 2019), and the rate of giant spikes increased with age up to ∼6 giant spikes/hour at 14 weeks of age (Figure 3E).

**Figure 3.**
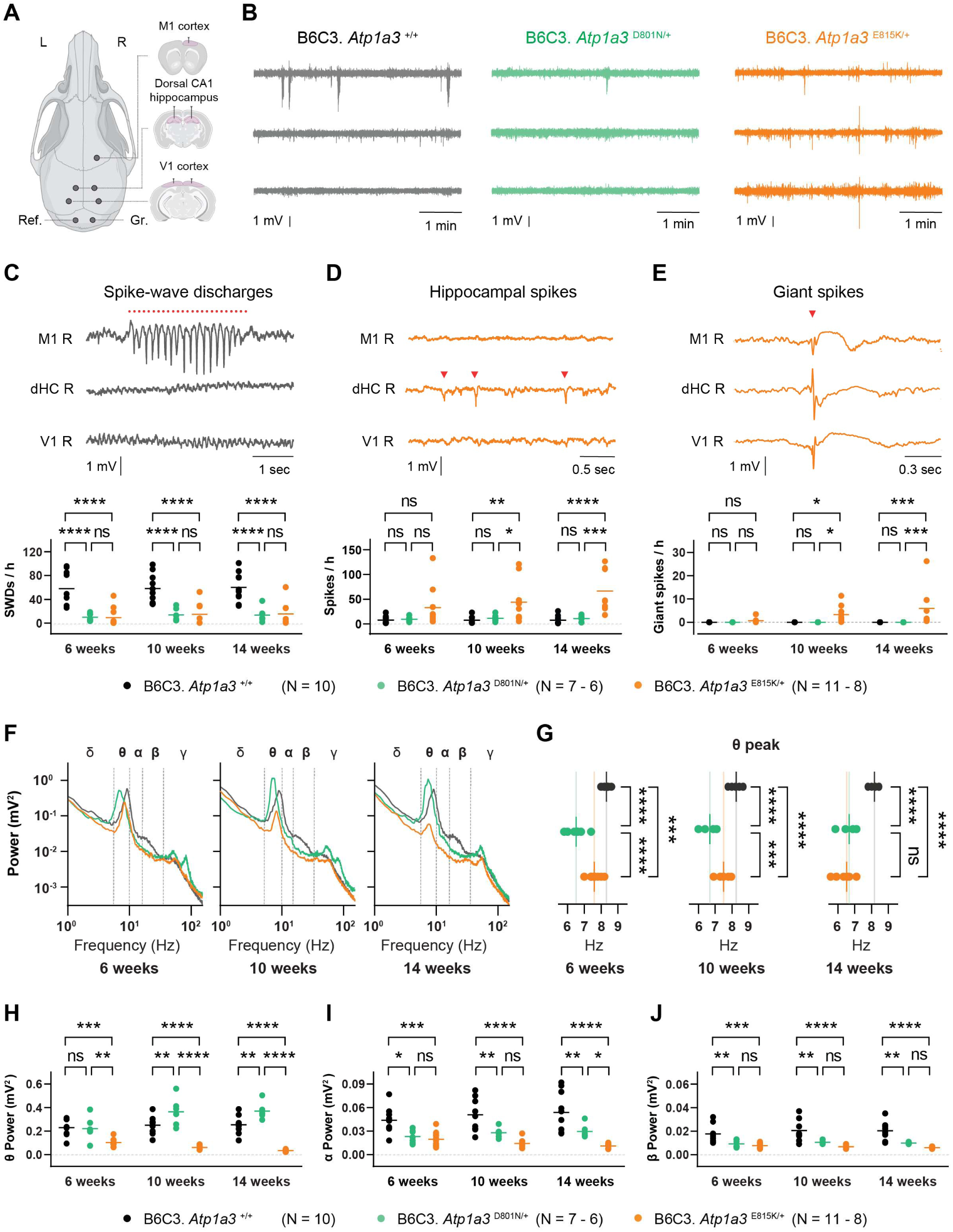
B6C3 AHC mice show altered hippocampal and cortical network activity. (A) Schematic representation of LFP recording electrodes placed in motor (M1) and visual (V1) cortex and dorsal CA1 hippocampus. Recordings were performed for 3 consecutive days per time point. Electrodes for reference (Ref.) and ground (Gr.) were placed in cerebellum. (B) Example traces of 5-minute AC-LFP traces from the M1, dorsal CA1 hippocampus, and V1 (top-to-bottom) of wild type (black), B6C3.*Atp1a3* ^D801N/+^ (green) and B6C3.*Atp1a3* ^E815K/+^ (orange) mice at 14 weeks of age. (C) Example trace of spike-wave discharge (SWD) event in frontal M1 cortical channel recorded in wild type mice at 14 weeks of age. SWDs present as a prominent burst (highlighted by the red dotted line) of negative polarity spikes and a positive polarity wave, exclusively seen in the frontal M1 cortical channel with no corresponding activity in V1 cortical or dHC channels. The horizontal line for each group represents the average SWD frequency per hour. (D) Example traces of hippocampal spike events (red arrowheads) in the dorsal CA1 hippocampus of B6C3.*Atp1a3* ^E815K/+^ mice at 14 weeks of age. These concerned isolated hippocampal spikes that were not observed in M1 and V1 cortices. The horizontal line for each group represents the average spike frequency per hour. (E) Example traces of hippocampo-cortical giant spike observed in B6C3.*Atp1a3* ^E815K/+^ mice. A giant spike consists of a simultaneous large spike of varying polarity (indicated by the red arrowhead) seen in all channels that is followed by a slow negative-positive potential shift lasting > 0.3 seconds in the M1 cortical channel and accompanied by a slow positive deflection lasting up to 600 ms in dHC and V1 cortex. The horizontal line for each group represents the average giant spike frequency per hour. (F) Average V1 cortical power spectral densities (PSD) during active wakefulness of wild type (black), B6C3.*Atp1a3* ^D801N/+^ (green) and B6C3.*Atp1a3* ^E815K/+^ (orange) mice. (G) Leftward shift of θ frequency peak observed in B6C3 AHC mice from 6 weeks onward. The vertical line for each group represents the mean. (H) Absolute power at θ frequency from the V1 cortical PSD during active wakefulness. The horizontal line for each group represents the mean. (I) Absolute power at α frequency from the V1 cortical PSD during active wakefulness. The horizontal line for each group represents the mean. (J) Absolute power at β frequency from the V1 cortical PSD during active wakefulness. The horizontal line for each group represents the mean. Ref, reference electrode; Gr, ground electrode; M1, motor cortex; V1, visual cortex; dHC, dorsal hippocampus CA1; CA1, hippocampal cornu ammonis area 1; SWD, spike-wave discharge. Two-way ANOVA was corrected for multiple comparisons using Tukey method (C, D, E, H, I, J); One-way ANOVA was corrected for multiple comparisons using Tukey method (G). ns, not significant; * p ≤ 0.05; ** p ≤ 0.01; *** p ≤ 0.001; **** p ≤ 0.0001. The depicted top view of the mouse skull with electrode configuration and coronal brain views were created using BioRender.com (A). See also Figures S3 and S4.

In addition to the spiking activities, we observed rare events in the cortical and hippocampal DC-potential recordings that are characterized as spreading depolarizations (SD). In B6C3.*Atp1a3* ^D801N/+^ mice a single spontaneous SD event was identified at 6 weeks of age that was first observed in motor (M1) cortex and then appeared in visual (V1) cortex, without appearing in the dorsal hippocampus (dHC) even though the LFP signal was also suppressed in this brain structure (Figure S3A). The SD event was preceded by a burst of spiking activity in M1 cortex (Figure S3B). In contrast, ∼50% of B6C3.*Atp1a3* ^E815K/+^ mice exhibited spontaneous SD events (example traces in Figure S3C, E and G). The SDs were consistently first observed in hippocampus (99% bilaterally) and with a small delay also appeared in cortex (Figure S3C and E). The likelihood that hippocampal SDs also appeared in cortex increased with age (20% of SDs at 10 weeks; 54% of SDs at 14 weeks). When appearing in cortex, SD events were first observed in the V1 cortex (43% unilateral and 57% bilateral) and from there propagated in a caudal-to-rostral pattern to M1 cortex in 85% of SD events (Figure S3E). Hippocampal SDs in B6C3.*Atp1a3* ^E815K/+^ mice were preceded by an epileptiform burst (Racine scale score 1-3) (Racine, 1972) that was characterized by high-amplitude spikes in hippocampal LFP synchronous with lower-amplitude spikes in cortical LFP (Figure S3D and F). The duration of the hippocampal burst activity preceding the SDs was not different for SDs that did or did not appear in cortex (Figure S3H). However, SD events with the appearance in the cortex exhibited higher burst spiking frequency as well as prolonged time to 50% SD recovery and a higher amplitude in hippocampus (Figure S3H).

To get a more detailed perspective of the brain activity changes, visual cortex AC-LFP recordings were used to analyze the power spectral density (PSD). The PSD from mouse LFP can be categorized into five frequency bands (i.e., δ (1-5 Hz), θ (5-10 Hz), α (10-13 Hz), β (13-30 Hz), and γ (30-100 Hz) bands) (Figure S4A), which in animals and humans are associated with various neurological activities and behaviors and when altered may provide insight in disease mechanisms (Drinkenburg et al., 2015; Tivadar & Murray, 2019). We first focused on the θ band because changes could be indicative of abnormal cognitive and motor function as well as epilepsy (Jansen et al., 2021; Kropotov, 2009; Luo et al., 2024; Perez et al., 2024; Tan et al., 2024). During periods of active wakefulness, characterized by faster frequencies of brain activity and high levels of θ activity (V1 cortical LFP example in Figure S4B), PSD analysis revealed a leftward shift in the θ frequency peak in both B6C3 AHC mice (Figure 3F and G). While the θ frequency peak in B6C3.*Atp1a3* ^D801N/+^ mice was consistently left-shifted, the θ frequency peak in B6C3.*Atp1a3* ^E815K/+^ mice gradually declined with age to the frequency observed in B6C3.*Atp1a3* ^D801N/+^ mice (Figure 3F and G). Moreover, the θ power gradually increased in B6C3.*Atp1a3* ^D801N/+^ while it decreased in B6C3.*Atp1a3* ^E815K/+^mice (Figure 3H). In addition, we observed significant decreases in α and β power in both B6C3 AHC mice, but these did not overtly change with age (Figure 3I and J). During periods of quiet wakefulness (lacking prominent θ activity, V1 cortical LFP example in Figure S4C), however, only B6C3.*Atp1a3* ^E815K/+^ mice exhibited a gradual reduction in power across the θ, α and β frequencies (Figure S4D, E, F and G).

Our EEG recordings show altered brain activity, with a reduction in the θ frequency peak in both B6C3 AHC mice, potentially reflecting an impairment in inhibitory neurotransmission. However, the distinct spectral power changes observed for the θ band for the two strains of B6C3 AHC mice suggests differences in synaptic strength with an increase in B6C3.*Atp1a3* ^D801N/+^ (increased θ power) and a decrease in B6C3.*Atp1a3* ^E815K/+^ (reduced θ power) mice. The observation of AHC variant specific brain activity changes may hint at potentially different mechanisms contributing to the development of distinct behavioral changes observed in B6C3 AHC mice. Importantly, B6C3.*Atp1a3* ^E815K/+^ mice revealed strikingly prominent electrophysiological changes (e.g., increased number of hippocampal spikes, occurrence of hippocampal-cortical giant spikes, SD events) that have previously been associated with an increased risk of epilepsy development (Buzsáki, 2015; Loonen et al., 2019; Motelow & Blumenfeld, 2009; Tamim et al., 2021; Zhen et al., 2021) and thereby, align with the increased seizure susceptibility observed in AHC patients. The epileptiform features and SD events observed in B6C3 AHC mice were not associated with profound behavioral seizure activity, nor can they explain the higher mortality of B6C3.*Atp1a3* ^D801N/+^ mice. However, our brain activity recordings were only performed over 3 days per time point in each mouse and none of the mice who died during the experiment (1 out of 7 D801N mice, 3 out of 11 E815K mice) happened to be in an EEG/video recording cage at the time of death. Neurophysiological abnormalities linked to mortality may require continuous and long-term EEG recordings to capture possible alterations that occur just prior to the unexpected death of B6C3 AHC mice.

### Neurological changes in AHC mice affect neuronal health and neuroinflammation

To investigate whether the neurological defects in B6C3 AHC mice are accompanied by molecular and cellular alterations, we next performed transcriptome analysis on different brain regions including the cortex, hippocampus, brainstem and cerebellum of wild type, B6C3.*Atp1a3* ^D801N/+^ and B6C3.*Atp1a3* ^E815K/+^ mice (Figure 4A). Aiming to detect potential molecular changes that are evoked upon neurological deficits, we analyzed adult B6C3 AHC mice (10 weeks) by which age mice are already susceptible to stress-induced paroxysmal spells, and exhibit motor defects and neurophysiological alterations, but prior to the overt onset of spontaneous deaths.

**Figure 4.**
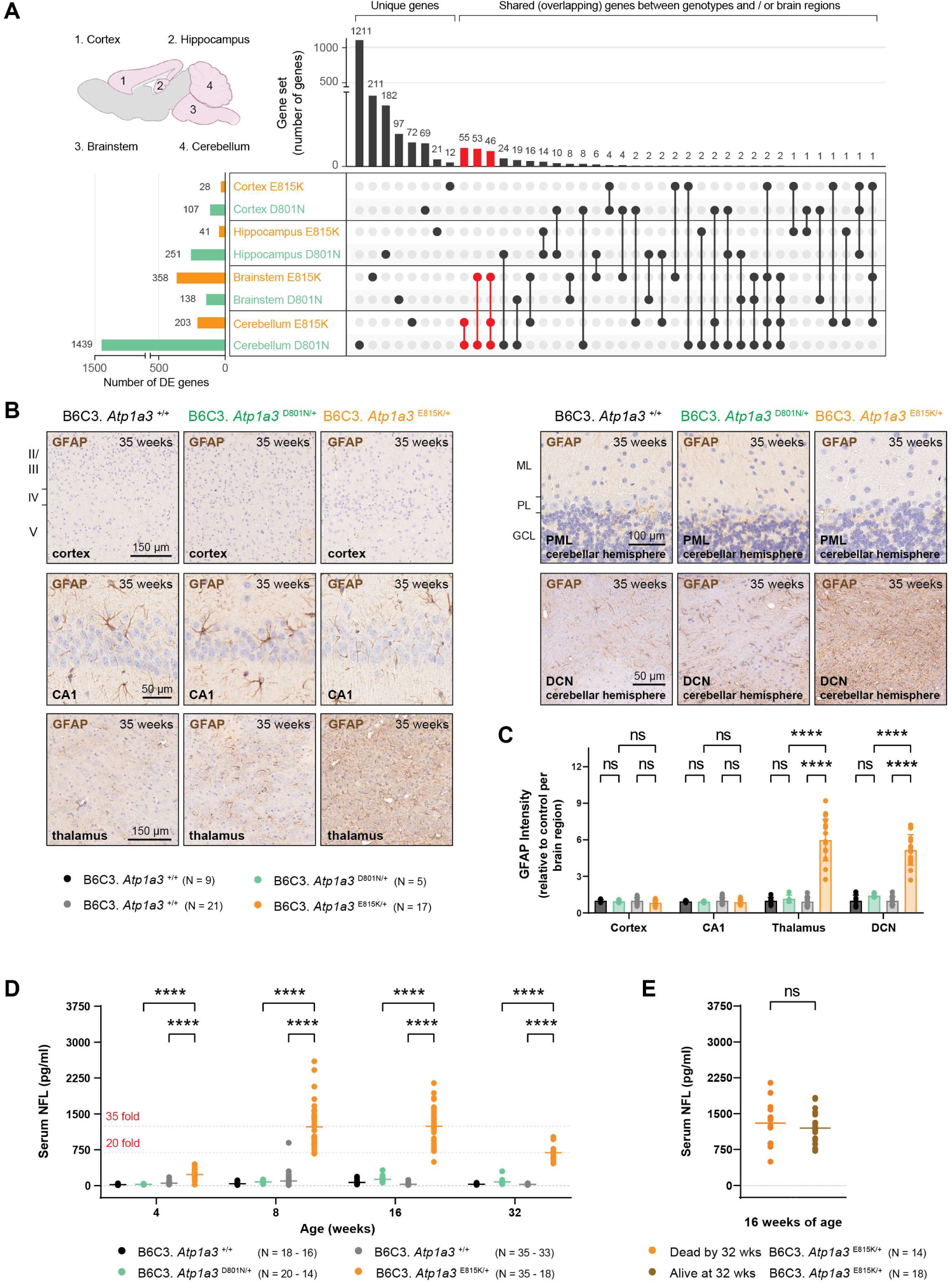
Gene expression analysis of different brain regions in B6C3 AHC mice. (A) UpSet plot of differentially expressed (DE) genes (adj. *p* ≤ 0.05) comparing AHC mutations and brain regions in mice. The anatomical locations of the analyzed mouse brain regions are shown in the upper, left corner. The total number of differentially expressed (adj. *p* ≤ 0.05) genes for each brain region of B6C3.*Atp1a3* ^D801N/+^ (D801N, green) and B6C3.*Atp1a3* ^E815K/+^ (E815K, orange) mice are shown in the lower left corner. The number of unique genes and the intersection of genes are shown on the right-hand side. Notable gene intersections between genotypes and brain region are highlighted in red. (B) Immunohistochemistry using antibodies to glial fibrillary acidic protein (GFAP, astrocytes, brown) on parasagittal brain sections from 35-week-old wild type, B6C3.*Atp1a3* ^D801N/+^ and B6C3.*Atp1a3* ^E815K/+^ mice. Sections were counterstained with hematoxylin and higher magnifications of individual brain regions are shown. (C) Relative intensity of GFAP of wild type, B6C3.*Atp1a3* ^D801N/+^ and B6C3.*Atp1a3* ^E815K/+^ mice. Levels are relative to that of controls (wild type) from each cross and brain region. Data represent mean ± SD. Wild type littermate controls for each mutant strain are shown (black and grey). (D) Longitudinal serum analysis of neurofilament light chain (NFL) of B6C3 AHC mice. The horizontal line for each group represents the mean. Wild type littermate controls for each mutant strain are shown (black and grey). (E) Comparison of serum NFL levels between B6C3.*Atp1a3* ^E815K/+^ mice that were either deceased at a later age (orange) or remained alive (brown). The horizontal line for each group represents the mean. Scale bars: 150 μm (cortex), 50 μm (CA1), 150 μm (thalamus), 100 μm (cerebellar hemisphere, PML) and 50 μm (cerebellar hemisphere, DCN). DE, differential gene expression; CA1, hippocampal cornu ammonis area 1; ML, molecular cell layer; PL, Purkinje cell layer; GCL, granule cell layer; PML, paramedian lobe; and DCN, deep cerebellar nuclei. Two-way ANOVA was corrected for multiple comparisons using Tukey method (C, D); Student’s t-test (E). ns, not significant; **** p ≤ 0.0001. The depicted mouse brain with specific brain regions was created with BioRender.com (A). See also Figures S5, S6 and S7.

Relative to wild type mice, differential gene expression (DE) analysis revealed several significant (adj. *p* ≤ 0.05) gene expression changes in B6C3 AHC mice and expression for a subset of genes was confirmed by quantitative RT-PCR (TaqMan) (Supplementary file 5, Figure S5A and B). Similar to the ATP1A3 protein expression (Figure S1F and G), no significant changes in *Atp1a3* gene expression were observed in B6C3 AHC mice (Supplementary file 5). To get a broader perspective of the gene expression changes, we visualized the intersection of the differentially expressed genes between AHC linked mutations and brain regions using an ‘Upset plot’. Except for the brainstem, B6C3.*Atp1a3* ^D801N/+^ mice generally showed a higher number of differentially expressed genes than B6C3.*Atp1a3* ^E815K/+^ mice (Figure 4A). When comparing the differentially expressed genes between D801N and E815K for each brain region, no significant gene overlap (hypergeometric density, Benjamini-Hochberg) was observed in the cortex, hippocampus and brainstem, which parallels the Upset plot that revealed only a few shared genes between B6C3.*Atp1a3* ^D801N/+^ and B6C3.*Atp1a3* ^E815K/+^ mice (Figure 4A). However, a significant (*p* = 3.6558 × 10^-3^, hypergeometric density, Benjamini-Hochberg) gene overlap was detected when comparing the cerebellar genes of B6C3.*Atp1a3* ^D801N/+^ (7.5% of D801N genes overlap with E815K) and B6C3.*Atp1a3* ^E815K/+^ mice (53% of E815K genes overlap with D801N) (Figure 4A). Although not significant (*p* = 0.06313), a moderate gene overlap was also observed between genes of the cerebellum and brainstem in B6C3.*Atp1a3* ^E815K/+^ mice (Figure 4A), highlighting that while most of the gene expression appeared to be unique in B6C3 AHC mice, at least a subset of the differentially expressed genes are affected in multiple brain regions by either mutation (red, Figure 4A).

To identify pathways that might be linked to the cerebellar and brainstem gene expression changes, we performed KEGG and IPA pathway analysis on the differentially expressed (adj. *p* ≤ 0.05) genes of B6C3.*Atp1a3* ^D801N/+^ and B6C3.*Atp1a3* ^E815K/+^ mice. Strikingly, immune response and inflammation related pathways were highly enriched and represented most of the top pathways in the cerebellum of B6C3.*Atp1a3* ^E815K/+^ mice (pathways are highlighted in red, Figure S5C and D). However, immune system related pathways showed a noticeable lower enrichment in the cerebellum of B6C3.*Atp1a3* ^D801N/+^ mice and mirrored the pathway enrichment in the brainstem of B6C3 AHC mice (Figure S5C, D, E and F).

The comparison with neuroinflammatory genes that were previously identified in CNS tissues of mouse models with various neurological and neurodegenerative conditions (Supplementary file 7) (Chiu et al., 2013; Holtman et al., 2015; Ishimura et al., 2016; Orre et al., 2014) revealed that approximately 18% and 27% of the differentially expressed cerebellar genes (adj. *p* ≤ 0.05) were neuroinflammation genes in B6C3.*Atp1a3* ^D801N/+^ and B6C3.*Atp1a3* ^E815K/+^ mice, respectively (Figure S6A and B). Interestingly, expression of these neuroinflammatory genes was significantly increased (fold change) in both B6C3 AHC models (Figure S6C). Nevertheless, B6C3.*Atp1a3* ^E815K/+^ showed a higher upregulation (fold change) of neuroinflammatory genes compared to B6C3.*Atp1a3* ^D801N/+^ mice (Figure S6C). Moreover, only 7% of the inflammation genes showed an increase in gene expression greater than 2-fold in B6C3.*Atp1a3* ^D801N/+^ while 35% of neuroinflammatory genes showed an upregulation greater than 2-fold in B6C3.*Atp1a3* ^E815K/+^ mice. In addition, expression of neuroinflammatory genes was also significantly increased in the brainstem of B6C3.*Atp1a3* ^E815K/+^ mice (Figure S6D). Thus, similar to the pathway enrichment observations (Figure S5C, D, E and F), the magnitude and/or activation of neuroinflammatory related gene expression changes may differ between B6C3 AHC mice and brain regions. Because the previously identified neuroinflammatory genes included those of microglia and astrocyte origin (Supplementary file 7), these data suggest that mutations in the neuron specific ATP1A3 may also elicit cellular responses in cell types beyond ATP1A3 expressing neurons.

Since the transcriptome analysis revealed inflammation related gene expression changes, we performed immunostaining using antibodies against common neuroinflammation markers including GFAP (glial fibrillary acidic protein, astrocytes) and Iba1 (ionized calcium-binding adapter molecule 1, microglia) (Amalia, 2021; Cao et al., 2021; Ishimura et al., 2016; Y. Wang et al., 2023; Young et al., 2013). We did not observe overt changes in GFAP or Iba1 in the cortex and hippocampus of B6C3 AHC mice (Figure 4B and C, Figure S7A and B), which aligns with the lack of enrichment of immune related pathways in those brain regions. However, GFAP and Iba1 staining were strongly increased in the thalamus and deep cerebellar nuclei (DCN) in the cerebellum of B6C3.*Atp1a3* ^E815K/+^ mice (Figure 4B and C, Figure S7A and B). Moreover, B6C3.*Atp1a3* ^D801N/+^ mice only showed a significant increase in Iba1 reactivity in the deep cerebellar nuclei of the cerebellum whereas Iba1 and GFAP levels in the other regions of the brain remained similar to wild type mice (Figure 4B and C, Figure S7A and B). Interestingly, reactivity of these neuroinflammation markers was significantly stronger in B6C3.*Atp1a3* ^E815K/+^ compared to B6C3.*Atp1a3* ^D801N/+^ mice (Figure 4C, Figure S7B) and paralleled the directionally of the transcriptomic inflammatory pathways observed in B6C3 AHC mice (Figure S5C and D, Figure S6A, B, C, and D).

Furthermore, we longitudinally measured serum levels of neurofilament light chain (NFL) as a means to monitor ‘brain and neuronal health’ in B6C3 AHC mice. NFL is a neuron specific cytoskeletal protein, which has gained great attention as an unspecific biomarker to assess disease progression and pathology, as well as treatment response in animal models and humans. Increased NFL levels (e.g., in the cerebrospinal fluid, serum and plasma) have been observed upon neuronal and axonal damage, brain lesions, astrogliosis and immune cell extravasation (neuroinflammation), and impairment of the blood-brain-barrier integrity (Barro et al., 2020; Bavato et al., 2024; Freedman et al., 2024; Jing et al., 2024; Uher et al., 2021; Van Den Bosch et al., 2022; Yuan & Nixon, 2021). In B6C3.*Atp1a3* ^D801N/+^ mice, serum NFL levels remained comparably low to those observed in wild type mice and only showed a statistically significant (*p*-value = 0.0218) increase of ∼1.85 fold at 16 weeks of age (Figure 4D). In contrast, serum NFL levels progressively increased and leveled to a ∼20-fold increase in B6C3.*Atp1a3* ^E815K/+^ mice (Figure 4D). In the case of other neurological disorders, increased NFL levels negatively correlated with survival while in others, the levels coincided with specific pathological cellular or functional alterations of the brain (Bacioglu et al., 2016; Bavato et al., 2024; Gravesteijn et al., 2019; Jung & Damoiseaux, 2023; Nguyen et al., 2022). In B6C3 AHC mice, serum levels of NFL did not predict or correlate with the observed survival defects (Figure 4D and E, Figure 1A and B). Regardless, the increase in NFL levels coincides with functional (e.g., motor and neurophysiological function) and histological (e.g., neuroinflammation) differences that are a more severe in B6C3.*Atp1a3* ^E815K/+^ compared to B6C3.*Atp1a3* ^D801N/+^ mice. Thus, changes in NFL levels may not holistically capture disease severity in B6C3 AHC mice that is reflected or assumed by their mortality, but perhaps, points to the differential severity of specific functional and/or morphological changes in the brain, which are linked to certain AHC variants.

## Discussion

AHC patients with the p.D801N or p.E815K variant exhibit paroxysmal spells by the age of 18 months. When comparing the two AHC variants, p.E815K patients present an earlier onset, a further delay in unsupported sitting and walking, and more severe cognitive impairment, dystonia and seizures (Capuano et al., 2020; Ford et al., 2023; Panagiotakaki et al., 2015; Viollet et al., 2015). In addition, patients with the p.E815K variant have a significantly higher occurrence (∼3 times) of status epilepticus (prolonged or repetitive seizures), suggesting that a higher pathogenicity may be linked to the p.E815K variant relative to other AHC variants (Capuano et al., 2020; Ford et al., 2023; Panagiotakaki et al., 2015; Viollet et al., 2015). A similar notion of p.E8151K causing more severe neurological abnormalities was suggested for the previously developed B6J AHC mice (D801N and E815K) with the B6J E815K mice revealing rotarod defects and increased freezing behavior compared to B6J D801N mice (Helseth et al., 2018; Hunanyan et al., 2014). However, comparing and defining similarities or dissimilarities of AHC linked mutations in B6J AHC mice remains difficult, at least in part, because of challenges that arose from the severe mortality and/or differences in experimental testing paradigms.

B6C3.*Atp1a3* ^E815K/+^ mice showed overt electrophysiological abnormalities (e.g. spikes and spontaneous SDs) compared to B6C3.*Atp1a3* ^D801N/+^ mice, fitting with the higher susceptibility of epilepsy observed in p.E815K patients. Notably, B6C3 wild type mice exhibited a higher incidence of SWDs compared to B6C3 AHC mice. The incidence of SWDs, with a wider variation, in B6C3 mice is consistent with the introduction of C3H/HeJ (F1 hybrid with C57BL/6J) compared to the parental C3H/HeJ strain (Frankel et al., 2005), highlighting the complex interaction of genetic factors between C3H/HeJ and C57BL/6J that contribute to presence of SWDs. However, why B6C3 AHC mice show fewer SWDs compared to B6C3 wild type mice remains unclear. The reduced synaptic strength, as shown by the EEG power analysis, may affect activity to sustain a high frequency of SWDs while being sufficient to increase the occurrence of hippocampal and giant spikes in B6C3.*Atp1a3* ^E815K/+^ mice. In addition, motor (e.g., rotarod performance) and molecular defects (e.g., NFL, neuroinflammation) were also more affected in B6C3.*Atp1a3* ^E815K/+^ mice. While these deficits align with the severe impact of the p.E815K variant, spontaneous lethality and hypothermia induced paroxysmal episodes were instead more prominent in B6C3.*Atp1a3* ^D801N/+^ mice. Furthermore, differential responses were also observed in the open field testing (e.g., horizontal and vertical activity) and during electrophysiology recordings (e.g., θ power) of B6C3 AHC mice. These data imply that tractable and distinct alterations for specific AHC variants can be detected in mice and perhaps, phenotypic severity may not merely be a function of a specific genotype (e.g., p.E815K being the most severe variant) but may also depend on the specific defect that is investigated. Recently, the third most common AHC variant (p.G947R) was introduced in B6J mice, which also causes electrophysiological defects and stress induced dystonia (Hawkins et al., 2024). Interestingly, B6N D801Y mice, which carry an ATP1A3 variant linked to RDP/ AHC and B6J G947R mice showed an increase in rotarod performance that is in contrast to the reduced rotarod performance observed in B6C3 D801N and E815K mice (Y. B. Liu et al., 2024; Ozelius, 2004; Viollet et al., 2015). Although these strains differ in their genetic background and to a certain extent could limit a fair comparison, these data may still hint to specific changes that are mediated in an AHC variant dependent manner. Thus, continuing efforts to interrogate and systemically compare alterations of mice that harbor different pathogenic AHC variants may provide viable approaches to identify and delineate faulty neuronal circuits regulating dystonia, seizure and/or motor activity defects in AHC.

Even though ATP1A3 expression is restricted to neurons in the brain (Dobretsov et al., 2019; Gokce et al., 2016; Jiao et al., 2022; Smith et al., 2021), adding to the investigation of neuronal function is perhaps the interaction with non-neuronal cells of the brain. The analysis of brains of B6C3 AHC mice revealed expression and activation of inflammatory markers in different brain regions. Whether these neuro- inflammatory changes have functional implications is unclear and will require further investigation. In the case of neurological disorders, neuroinflammation can be protective or deleterious when chronic and uncontrolled (Adamu et al., 2024; DiSabato et al., 2016). Numerous studies have highlighted the complex interaction between epilepsy and inflammation, in which neuroinflammation can be cause or consequence of seizures/epilepsy (Eyo et al., 2017; Foiadelli et al., 2023; Pracucci et al., 2021; Sanz et al., 2024; Suleymanova, 2021). Astrocytic and microglial ion pumps and channels can act as buffer systems of excessive extracellular ions during seizures and provide anti-seizure properties (Du et al., 2018; Eyo et al., 2014; Zhao et al., 2022). Nevertheless, cytokines that are released by astrocytes and microglia also have excitatory effects and may further burden brain excitability and/or cause neuronal excitotoxicity (Foiadelli et al., 2023; M. Liu et al., 2020; Sanz et al., 2024; Villasana-Salazar & Vezzani, 2023). Thus, anti-inflammatory drugs are also considered as potential treatment options when anti-convulsant drugs fail to show disease altering outcomes or patients have drug-resistant seizures/epilepsy (Pracucci et al., 2021; Radu et al., 2017; Sanz et al., 2024).

Deterioration in motor and intellectual function has been observed in some AHC patients and sometimes coincides with cerebral and cerebellar atrophy (Neville & Ninan, 2007; Paciorkowski et al., 2015; Sabouraud et al., 2019; Saito et al., 2010; Sasaki et al., 2014; Sweney et al., 2015). Irreversible atrophy is particularly more often noticed in p.E815K patients with status epilepticus (Grillo, 2015; M. Li et al., 2015; Sasaki et al., 2014, 2017). Because of the symptomatic heterogeneity and because progressive decline in neurological function is not always observed in AHC patients (Panagiotakaki et al., 2010; Pavone, Pappalardo, Mustafa, et al., 2022), it is still unclear whether pathogenic AHC variants cause progressive defects as part of the primary Na^+^/K^+^-ATPase defect, or perhaps, such defects are related to repeated paroxysmal episodes.

Interestingly, B6C3.*Atp1a3* ^E815K/+^ mice exhibited progressive deficits in motor function (e.g., rotarod performance), neurophysiology (e.g., SD, hippocampal spikes and giant spikes) and serum NFL levels in contrast to B6C3.*Atp1a3* ^D801N/+^ mice. Importantly, measurements for our studies were obtained from naïve and separate cohorts of mice with the aim to measure baseline phenotypic defects and minimize excessive stress that could affect functional readouts since B6J AHC mice were highly susceptible to stress (e.g., routine and experimental handling). While progressive defects can already be observed under baseline conditions, interrogating the neurological function of B6C3 AHC mice when intentionally also introducing stress paradigms (e.g., hypothermia-induced paroxysmal episodes, HIP) may provide opportunities to systematically test for the contribution of severe and repetitive paroxysmal episodes in AHC.

How different mutations in *ATP1A3* can lead to different phenotypic deficits and/or cellular responses is unclear. The D801N and E815K mutations abolish the catalytic activity of the mutant ATP1A3 protein *in vitro*. Interestingly, the pathogenic p.D801Y variant observed in RDP and mild AHC patients reduces ATP1A3 (α3) protein expression by ∼20% and α3-specific activity by ∼75% in the brain of B6N D801Y mice (Holm et al., 2016; Y. B. Liu et al., 2024; Ozelius, 2004; Viollet et al., 2015; K. Wang et al., 2023), which is less drastic compared to the impairment of α3 activity observed in B6C3 D801N mice and aligns with previous *in vitro* investigations (De Carvalho Aguiar et al., 2004; Heinzen et al., 2012). However, the mere functional impairment of ATP1A3 (loss of catalytic activity) does not correlate with the disease severity observed ATP1A3 patients (Arystarkhova et al., 2019; Lazarov et al., 2020). While both p.D801N and p.E815K ATP1A3 are catalytically inactive, previous studies found that the two variants differ in cation binding affinity and the p.E815K variant results in a severely affected loss-of-function protein, which may contribute, in part, to the severity observed in patients (Heinzen et al., 2012; Weigand et al., 2014). Nevertheless, the dominant nature of AHC variants suggests that additional mechanisms may also contribute to disease pathology and may include changes in protein folding, biosynthesis, and ATP1A3 membrane trafficking (Arystarkhova et al., 2019, 2021). Interestingly, recent studies demonstrated that human ATP1A3 interacts with RNA-binding proteins and proteins regulating translation, suggesting that ATP1A3 may potentially also play a role in RNA translation (Fujii et al., 2024). Whether pathogenic AHC variants alter such protein interactions and/or may even evoke aberrant interactions of the mutant protein, which has been observed for other genetic disorders such as Charcot-Marie-Tooth (CMT) disease, is unknown (Bervoets et al., 2019; Kalotay et al., 2023; Sun et al., 2021). However, these data and the increasing identification of non-canonical (non-catalytic) functions observed for other proteins open the path for investigations beyond the ‘canonical’ ion pump activity of ATP1A3 (Fujii et al., 2024; Huangyang & Simon, 2018; Jeffery, 2020; Pan et al., 2021; Snaebjornsson & Schulze, 2018).

Lastly, B6C3 AHC mice also provide alternate mouse models for translational and pre-clinical AHC research, and the strains have been donated to public mouse repositories (JAX/MMRRC). Fortunately, the introduction of hybrid vigor reduced early mortality and lethal episodes that occur during animal handling while these mouse models still develop clinically relevant phenotypes including spontaneous death, hypothermia induced paroxysmal episodes (HIP), and motor, neurophysiological and histological defects. For example, behavioral (e.g., rotarod and open field) and histological testing of mice present standardized, feasible and/or high throughput measurements. In comparison, HIP testing is generally more time consuming particularly when considering large scale efficacy studies. However, these experiments are compatible with camera systems to record the animal behavior during HIP recovery and in turn, simplify or at least add resolution when assessing the therapeutic response in mice. Moreover, biomarker discovery and development have become increasingly critical for both pre-clinical and early phase clinical trials to assess safety, efficacy or to molecularly demonstrate therapeutic engagement. The ability to longitudinally monitor disease and experimentally manipulate deficits in animal models enables future biomarker exploration when screening of such candidates in patient populations is challenging due to the limited number of patients, genetic, environmental or symptomatic heterogeneity. Although other ATP1A3 mouse models have been previously generated, expanding the existing repertoire (summarized by Ng et al., 2021) with the inclusion of these novel AHC models may not only assist in overcoming earlier experimental limitations but may also put forward new platforms for basic and translational ATP1A3 research.

---

**Figure S1.**
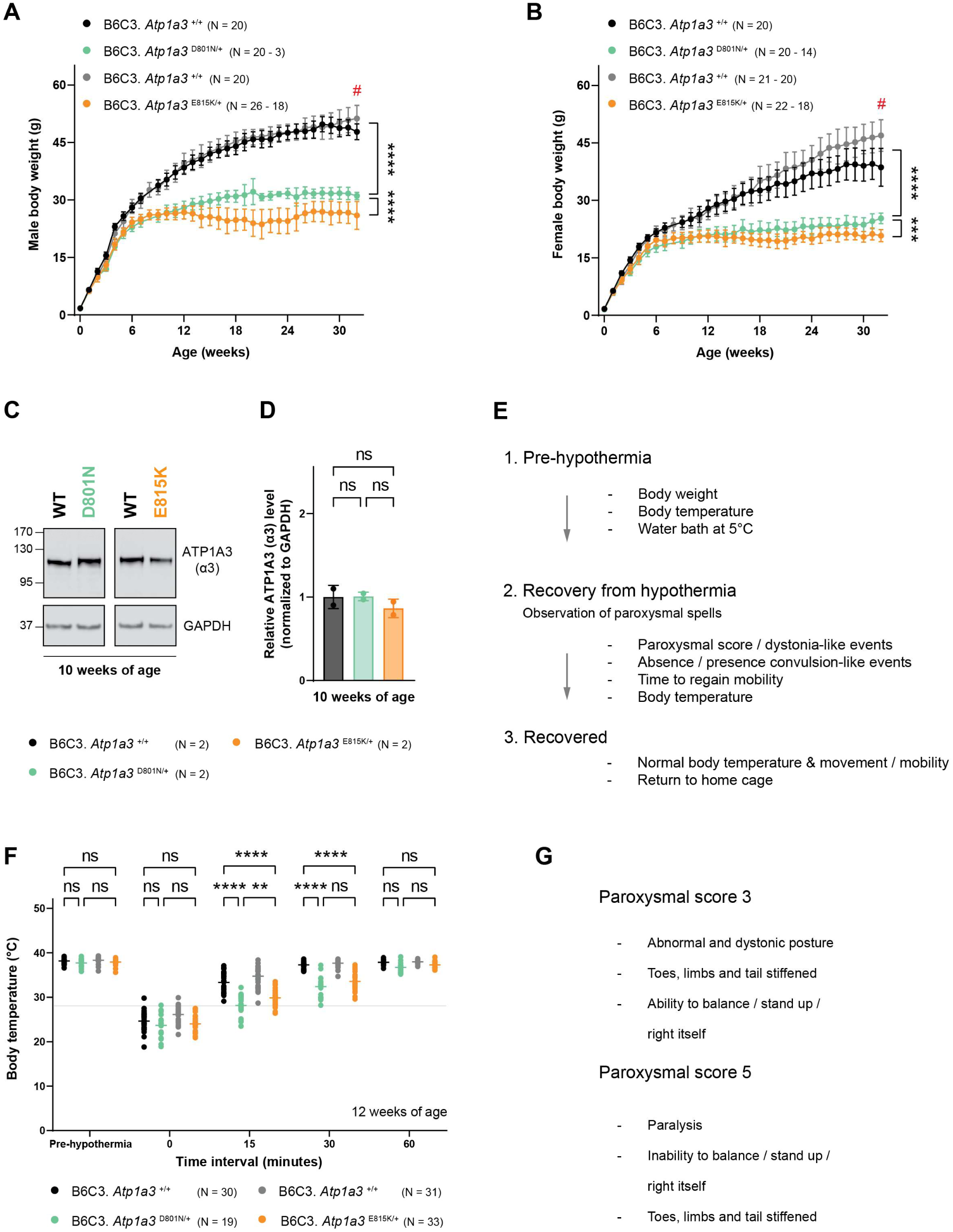
Hypothermia induced paroxysmal spells (HIP) in B6C3 AHC mice. (A) and (B) Body weight measurements of B6C3 AHC male (A) and female (B) mice. Wild type littermate controls for each mutant strain are shown (black and grey). **#** In the presence of a lowering body condition score, B6C3.*Atp1a3* ^E815K/+^ mice were provided with a diet gel. Because wild type (grey) and B6C3.*Atp1a3* ^E815K/+^ (orange) littermates are housed together, the body weight of wild type (grey) mice may exceed that of wild type (black) mice lacking access to this additional food source. Data represent mean ± SD. (C) Western blot analysis of ATP1A3 (α3) using hippocampal tissue samples from B6C3 AHC mice. GAPDH was used as loading control. (D) Relative expression of ATP1A3 (α3) subunit by western blotting of crude hippocampi homogenates. The ATP1A3 levels were normalized to levels of GAPDH. Protein levels are relative to those of wild type mice. Data represent mean + SD. (E) Experimental outline of HIP experiments. 1^st^ step: body temperature and body weight are measured. Mice are subjected to a 5°C water bath to induce hypothermia and the time spent in the water bath is relative to the body weight. 2^nd^ step: mice are monitored during the recovery of hypothermia for paroxysmal spells (dystonia- and conclusive-like events), regain of mobility and movement control and body temperature. 3^rd^ step: mice are returned to home cage once recovered (return to normal behavior and body temperature). (F) Body temperature monitoring of B6C3 AHC mice during HIP experiments. The body temperature is measured just prior to exposure to the water bath (pre-hypothermia), immediately after exposure to the water bath (0 minutes) and during recovery of hypothermia (15-, 30- and 60-minutes post water bath exposure). The horizontal line for each group represents the mean. Wild type littermate controls for each mutant strain are shown (black and grey). (G) Examples of paroxysmal scores. WT, wild type. ANOVA with repeated measurements was corrected for multiple comparisons using Tukey method (A, B). One-way ANOVA was corrected for multiple comparisons using Tukey method (C), Two-way ANOVA was corrected for multiple comparisons using Tukey method (F). ns, not significant; ** p ≤ 0.01; *** p ≤ 0.001; **** p ≤ 0.0001. Related to Figure 1.

**Figure S2.**
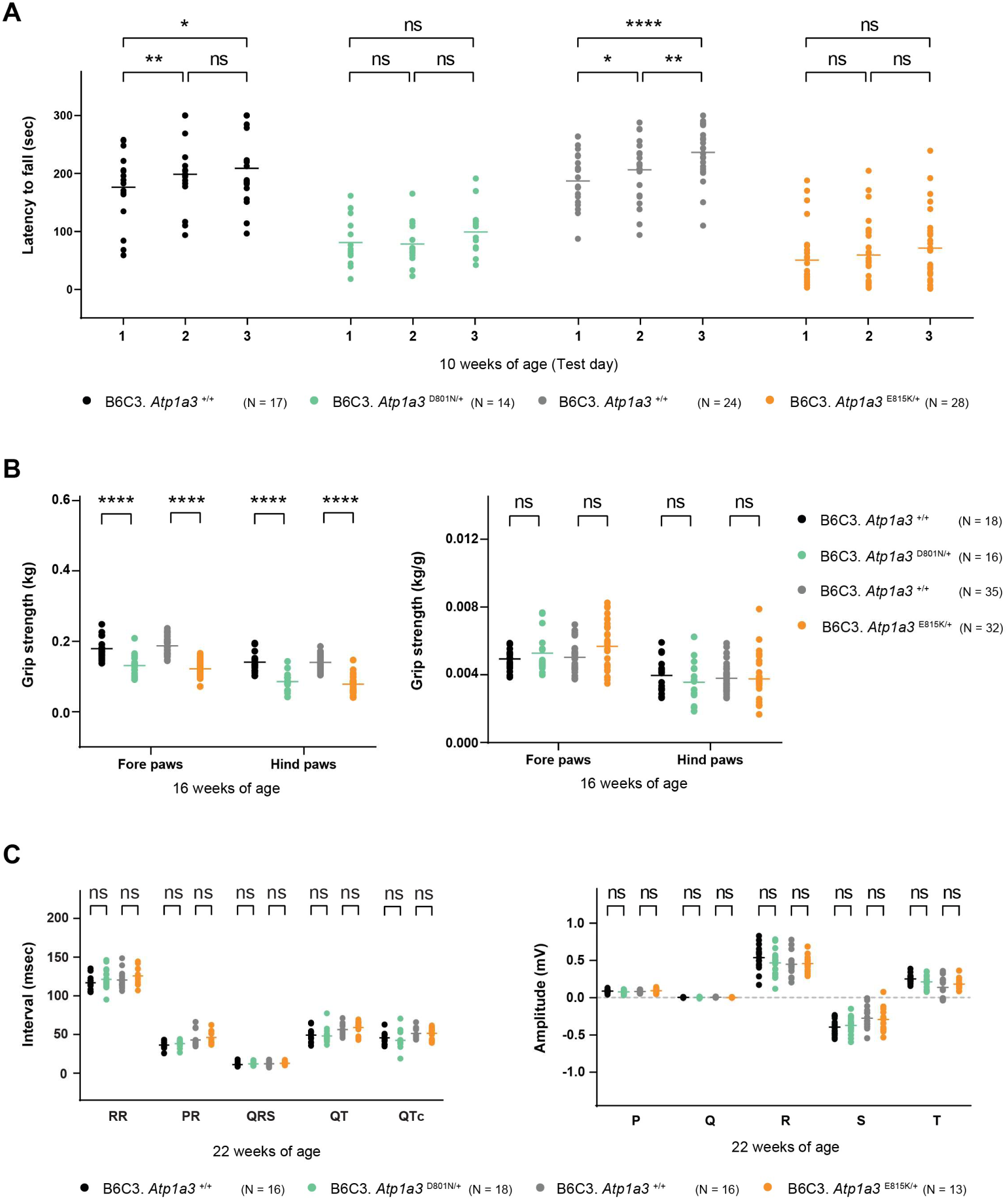
Behavioral alterations in B6C3 AHC mice. (A) Latency of B6C3 AHC mice to fall off the accelerating rotarod was measured over three consecutive testing days. The horizontal line for each group represents the mean. Wild type littermate controls for each mutant strain are shown (black and grey). (B) Grip strength was measured in B6C3 AHC mice. Data are shown without (left) and with (right) body weight normalization. The horizontal line for each group represents the mean. Wild type littermate controls for each mutant strain are shown (black and grey). (C) B6C3 AHC mice were subjected to electrocardiogram to measure the interval length and amplitude. The horizontal line for each group represents the mean. Wild type littermate controls for each mutant strain are shown (black and grey). Two-way ANOVA with repeated measurements was corrected for multiple comparisons using Tukey method (A), Two-way ANOVA was corrected for multiple comparisons using Tukey method (B, C). ns, not significant; * p ≤ 0.05; ** p ≤ 0.01; **** p ≤ 0.0001. Related to Figure 2.

**Figure S3.**
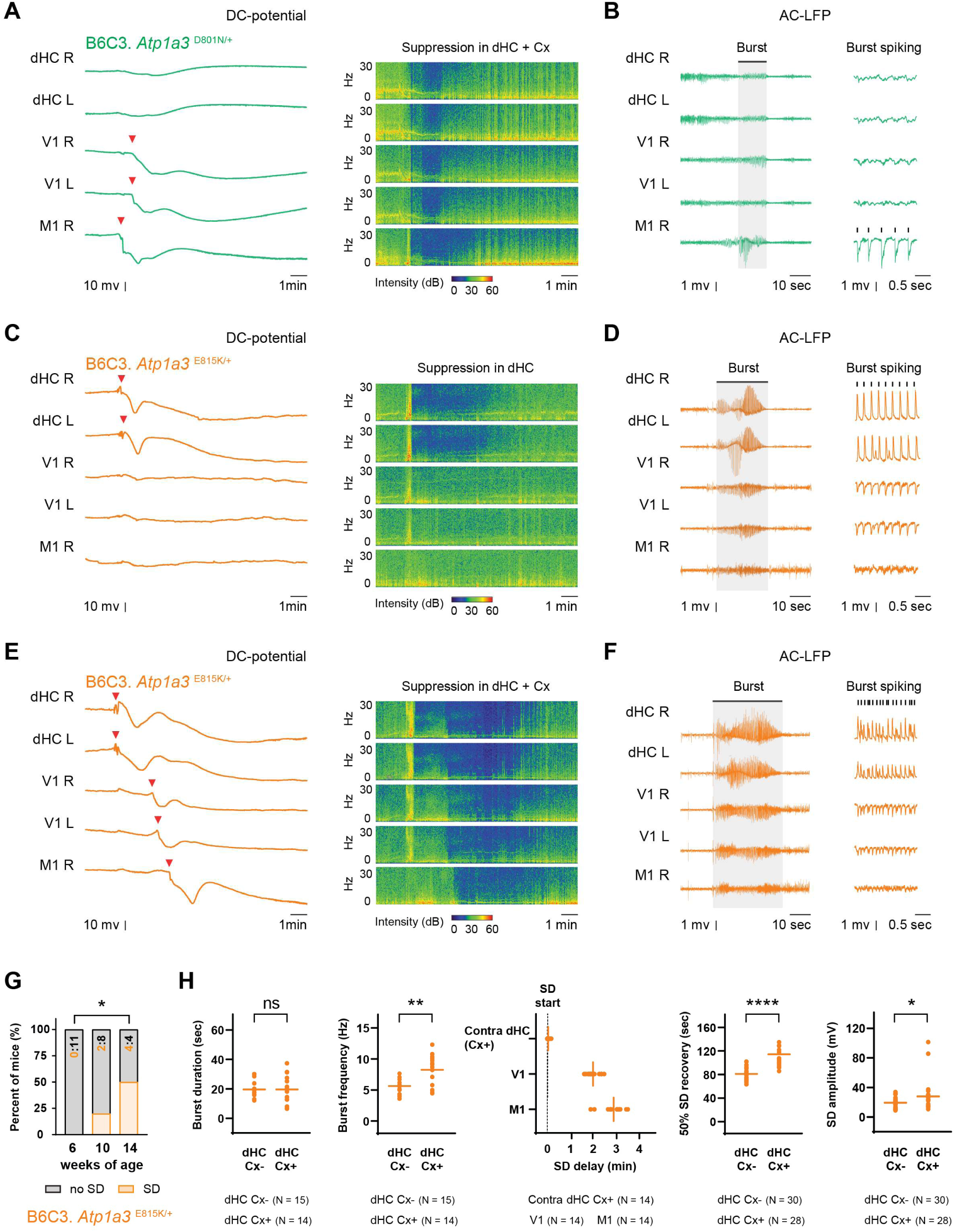
B6C3 AHC mice exhibit rare spontaneous spreading depolarization events. (A) Example of DC-potential trace of the spontaneously occurring SD event recorded in a B6C3.*Atp1a3* ^D801N/+^ mouse starting in M1 cortex with a spread to V1 cortex (red arrowhead indicates the > 5 mV negative DC-potential shift). Sonograms display SD-related suppression of LFP power across frequencies in hippocampal and cortical channels following the cortical SD event. (B) Example AC-LFP traces showing cortical (M1) burst activity (right insets showing burst spiking activity details) occurring prior to the SD event in a B6C3.*Atp1a3* ^D801N/+^ mouse. (C) Example DC-potential trace of a spontaneously occurring SD event (red arrowhead indicates the > 5 mV negative DC-potential shift) in a B6C3.*Atp1a3* ^E815K/+^ mouse with DC shifts occurring almost simultaneously in right (R) and left (L) dorsal hippocampi (dHC). Sonograms display SD-related suppression of LFP power across frequencies evident in dHC. (D) Example AC-LFP traces showing synchronously occurring burst activity, prior to the non-propagating SD event in panel C, with high-amplitude hippocampal and lower-amplitude cortical spikes observed bilaterally in dHC and cortical V1 channels, respectively. (E) Example DC-potential traces of a spontaneously occurring SD event (red arrowhead indicates the > 5 mV negative DC-potential shift) showing spread from hippocampus to cortex in a B6C3.*Atp1a3* ^E815K/+^ mouse with DC shifts occurring almost simultaneously in R and L dHC and sequentially propagating to V1 and M1 cortex. Sonograms display prolonged SD-related suppression of LFP power across frequencies in both hippocampal and cortical channels. (F) Example AC-LFP traces showing synchronously occurring burst activity, prior to the propagated SD event illustrated in (E), with high-amplitude hippocampal and lower-amplitude cortical spikes observed bilaterally in dHC and cortical V1 channels, respectively. (G) The occurrence of spontaneous SD events in B6C3.*Atp1a3* ^E815K/+^ mice. Data are shown as the fraction (percent) of mice without (grey) or with (orange) spontaneous SD events. The exact number of mice without (black font) and with (orange font) spontaneous SD events is shown in each bar. (H) Comparison of the dHC characteristics of non-propagating (Cx-) and propagating (Cx+) SD events in B6C3.*Atp1a3* ^E815K/+^ mice, including mean burst spiking frequency, burst duration, SD delay, 50% SD recovery time, and SD amplitude. The horizontal line for each group represents the mean. The sample size (N) reflects the number of non-propagating (Cx-) and propagating (Cx+) SD events observed in all B6C3.*Atp1a3* ^E815K/+^ mice. DC, direct current potentials; LFP, local field potential recording; R, right; L, left; dHC, dorsal hippocampus CA1; CA1, hippocampal cornu ammonis area 1; V1, visual cortex; M1, motor cortex; SD, spontaneous depolarization; Cx, cortex. Fisher’s exact test (G); Student’s t-test (H). ns, not significant; * *p* ≤ 0.05; ** *p* ≤ 0.01; **** *p* ≤ 0.0001. Related to Figure 3.

**Figure S4.**
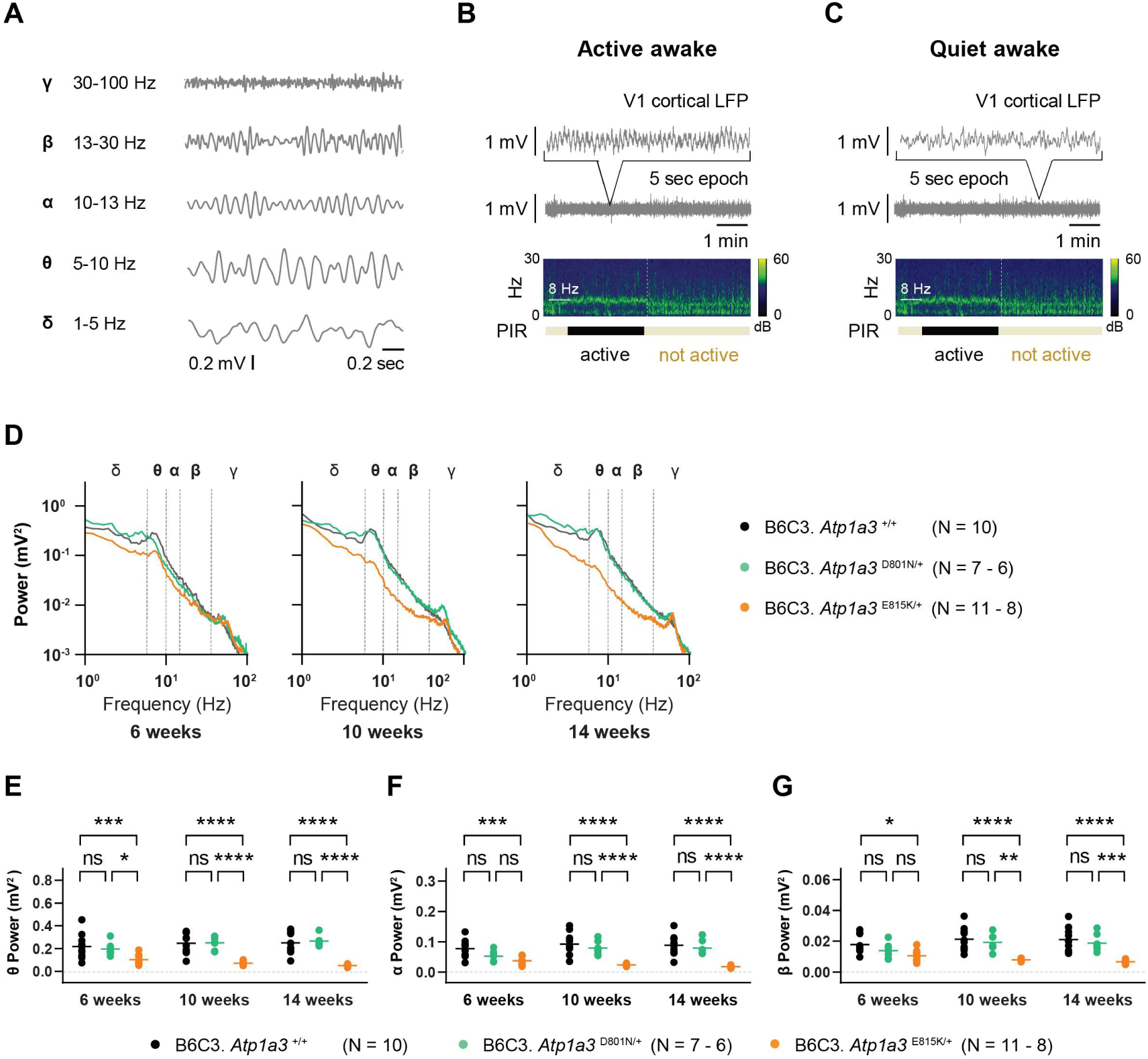
Neurophysiological alterations in the brains of B6C3 AHC mice during quiet wakefulness. (A) Examples of band-pass filtered δ, θ, α, β and γ signals. (B) Example trace of a 5-second V1 cortical LFP epoch when the mouse is active awake (showing also the transition of the active awake to an inactive state). During active wakefulness a prominent 8-Hz θ rhythm is observed in the sonogram and motor activity is present, as detected by a passive infrared motion sensor (PIR). C) Example trace of a 5-second V1 cortical LFP epoch during quiet wakefulness, as defined by a lack motor activity and lack of prominent 8-Hz θ rhythm. (D) Average V1 cortical LFP power spectral densities during quiet wakefulness in wild type (black), B6C3.*Atp1a3* ^D801N/+^ (green) and B6C3.*Atp1a3* ^E815K/+^ (orange) mice. (D) Average V1 cortical LFP power spectral densities during quiet wakefulness in wild type (black), B6C3.*Atp1a3* ^D801N/+^ (green) and B6C3.*Atp1a3* ^E815K/+^ (orange) mice. (E) Absolute power at θ frequency from the V1 cortical LFP power spectra during quiet wakefulness. The horizontal line for each group represents the mean. (F) Absolute power at α frequency from the V1 cortical LFP power spectra during quiet wakefulness. The horizontal line for each group represents the mean. (G) Absolute power at β frequency from the V1 cortical LFP power spectra during quiet wakefulness. The horizontal line for each group represents the mean. V1, visual cortex; PIR, infrared motion sensor; LFP, local field potential recording. Two-way ANOVA was corrected for multiple comparisons using Tukey method (E, F, G). ns, not significant; * p ≤ 0.05; ** p ≤ 0.01; *** p ≤ 0.001; **** p ≤ 0.0001. Related to Figure 3.

**Figure S5.**
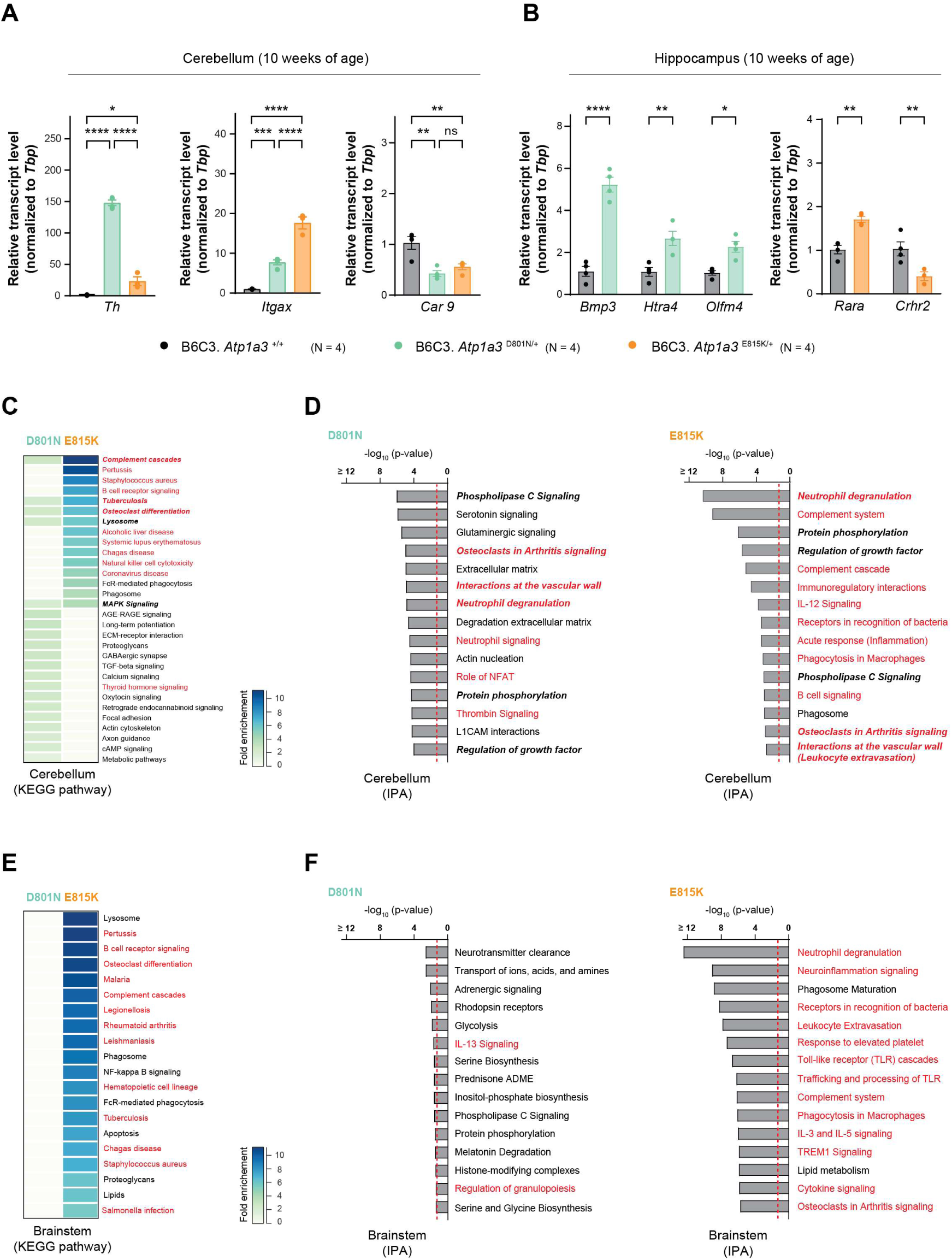
Pathway enrichment of differentially expressed genes in B6C3 AHC mice. (A) Differentially expressed (adj. *p* ≤ 0.05) cerebellar genes (*Th*, *Itgax* and *Car9*) identified from the transcriptome analysis of B6C3 AHC mice were analyzed via quantitative RT-PCR (TaqMan). Data were normalized to the transcript levels of TATA binding protein (*Tbp)* and the fold change in gene expression is relative to that of wild type mice. Data represent mean ± SEM. (B) Differentially expressed (adj. *p* ≤ 0.05) hippocampal genes (*Bmp3*, *Htra4, Olfm4, Rara* and *Crh2*) identified from the transcriptome analysis of B6C3 AHC mice were analyzed via quantitative RT-PCR (TaqMan). Data were normalized to the transcript levels of *Tbp* and the fold change in gene expression is relative to that of wild type mice. Data represent mean ± SEM. (C) KEGG pathway analysis (biological process) of differentially expressed (adj. *p* ≤ 0.05) cerebellar genes in B6C3.*Atp1a3* ^D801N/+^ (D801N, green) and B6C3.*Atp1a3* ^E815K/+^ (E815K, orange) mice. Significantly (*p* ≤ 0.05) enriched top pathways are shown. Pathways identified in both B6C3.*Atp1a3* ^D801N/+^ and B6C3.*Atp1a3* ^E815K/+^ mice are in bold italics. Pathways related to immune response are shown in red. (D) Ingenuity pathway analysis (IPA) of cerebellar genes in B6C3.*Atp1a3* ^D801N/+^ (D801N, green) and B6C3.*Atp1a3* ^E815K/+^ (E815K, orange) mice. Pathways identified in both B6C3.*Atp1a3* ^D801N/+^ and B6C3.*Atp1a3* ^E815K/+^ mice are in bold italics. Pathways related to immune response are shown in red. The red dashed line indicates the significance threshold (*p* = 0.05). (E) KEGG pathway analysis (biological process) of differentially expressed (adj. *p* ≤ 0.05) brainstem genes in B6C3.*Atp1a3* ^D801N/+^ (D801N, green) and B6C3.*Atp1a3* ^E815K/+^ (E815K, orange) mice. Significantly (*p* ≤ 0.05) enriched top pathways are shown for B6C3.*Atp1a3* ^E815K/+^ mice. No significant KEGG pathways were identified in B6C3.*Atp1a3* ^D801N/+^ mice in the brainstem. Pathways related to immune response are shown in red. (F) Ingenuity pathway analysis (IPA) of brainstem genes in B6C3.*Atp1a3* ^D801N/+^ (D801N, green) and B6C3.*Atp1a3* ^E815K/+^ (E815K, orange) mice. Pathways related to immune response are shown in red. The red dashed line indicates the significance threshold (*p* = 0.05). One-way ANOVA was corrected for multiple comparisons using Tukey method (A), Two-way ANOVA was corrected for multiple comparisons using Bonferroni method (B). ns, not significant; * p ≤ 0.05; ** p ≤ 0.01; *** p ≤ 0.001; **** p ≤ 0.0001. Related to Figure 4.

**Figure S6.**
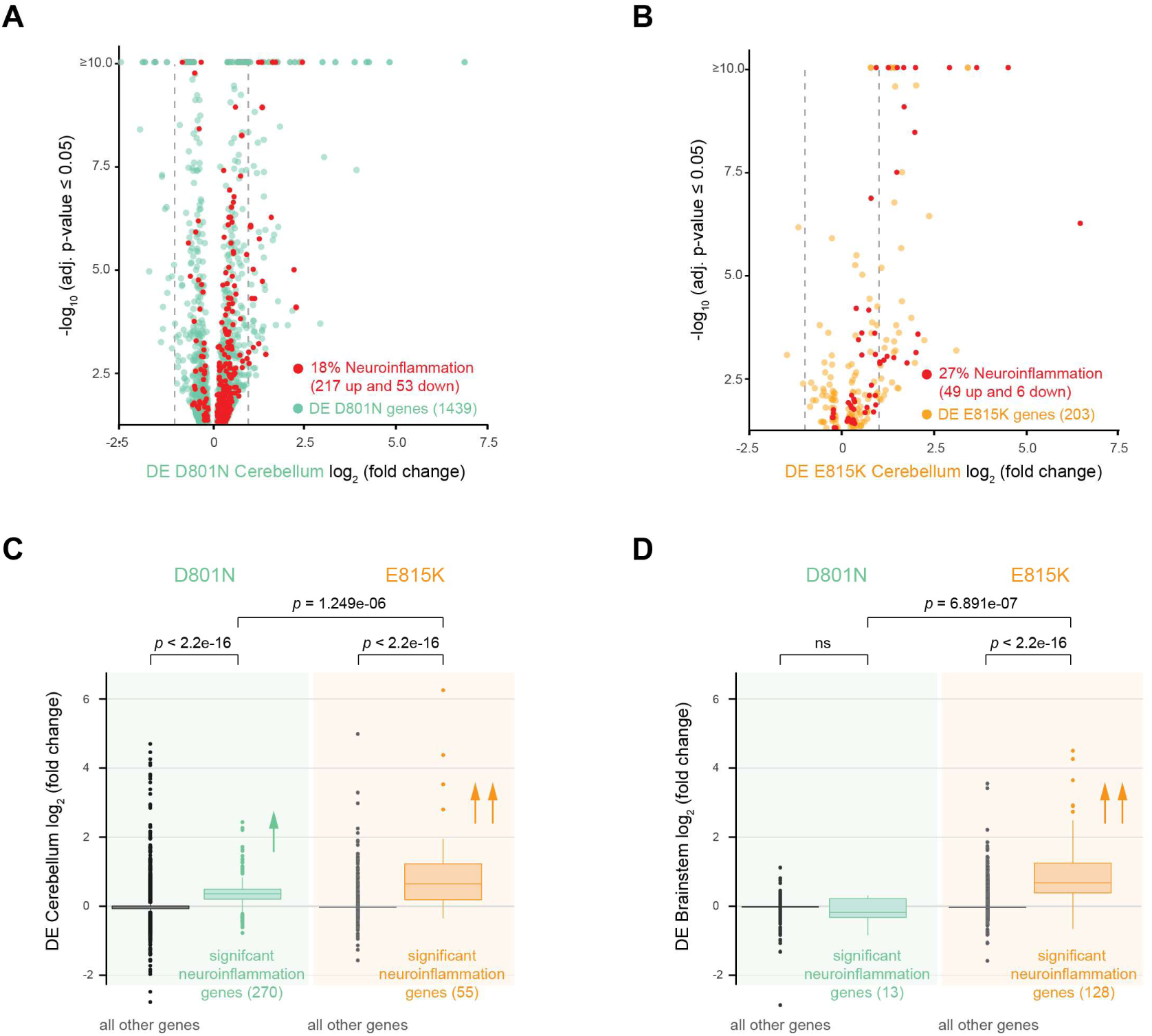
Gene expression of neuroinflammatory genes in the cerebellum of B6C3 AHC mice. (A) Analysis of differentially expressed cerebellar genes in B6C3.*Atp1a3* ^D801N/+^ mice (DE D801N cerebellum). Genes with significantly different expression (adj. *p* ≤ 0.05) in the cerebellum of D801N mice are shown in green (1439 genes). Of those cerebellar genes, neuroinflammation genes are highlighted in red (270 genes). Two-fold changes in gene expression are indicated by the grey dashed lines. (B) Analysis of differentially expressed cerebellar genes in B6C3.*Atp1a3* ^E815K/+^ mice (DE E815K cerebellum). Genes with significantly different expression (adj. *p* ≤ 0.05) in the cerebellum of E815K mice are shown in orange (203 genes). Of those cerebellar genes, neuroinflammation genes are highlighted in red (55 genes). Two-fold changes in gene expression are indicated by the grey dashed lines. (C) All cerebellar genes were compared to the significant (adj. p ≤ 0.05) differentially expressed neuroinflammatory genes of B6C3.*Atp1a3* ^D801N/+^ (D801N, 270 genes, green) and B6C3.*Atp1a3* ^E815K/+^ (E815K, 55 genes, orange). Upward direction of single arrow indicates significant increase in expression (positive fold change) of neuroinflammation genes in B6C3.*Atp1a3* ^D801N/+^ (D801N, green) compared to all other genes (black). Upward direction of double arrow indicates significant increase in expression (positive fold change) of neuroinflammation genes in B6C3.*Atp1a3* ^DE815K/+^ (E815K, orange) compared to all other genes (grey) and the inflammatory genes of B6C3.*Atp1a3* ^D801N/+^ (D801N, green) mice. (D) All brainstem genes were compared to the significant (adj. p ≤ 0.05) differentially expressed neuroinflammatory genes of B6C3.*Atp1a3* ^D801N/+^ (D801N, 13 genes, green) and B6C3.*Atp1a3* ^E815K/+^ (E815K, 128 genes, orange). Upward direction of double arrow indicates significant increase in expression (positive fold change) of neuroinflammation genes in B6C3.*Atp1a3* ^DE815K/+^ (E815K, orange) compared to all other genes (grey) and the inflammatory genes of B6C3.*Atp1a3* ^D801N/+^ (D801N, green) mice. DE, differential gene expression. Wilcoxon rank-sum test with continuity correction was used to determine statistical significance (C, D). Related to Figure 4.

**Figure S7.**
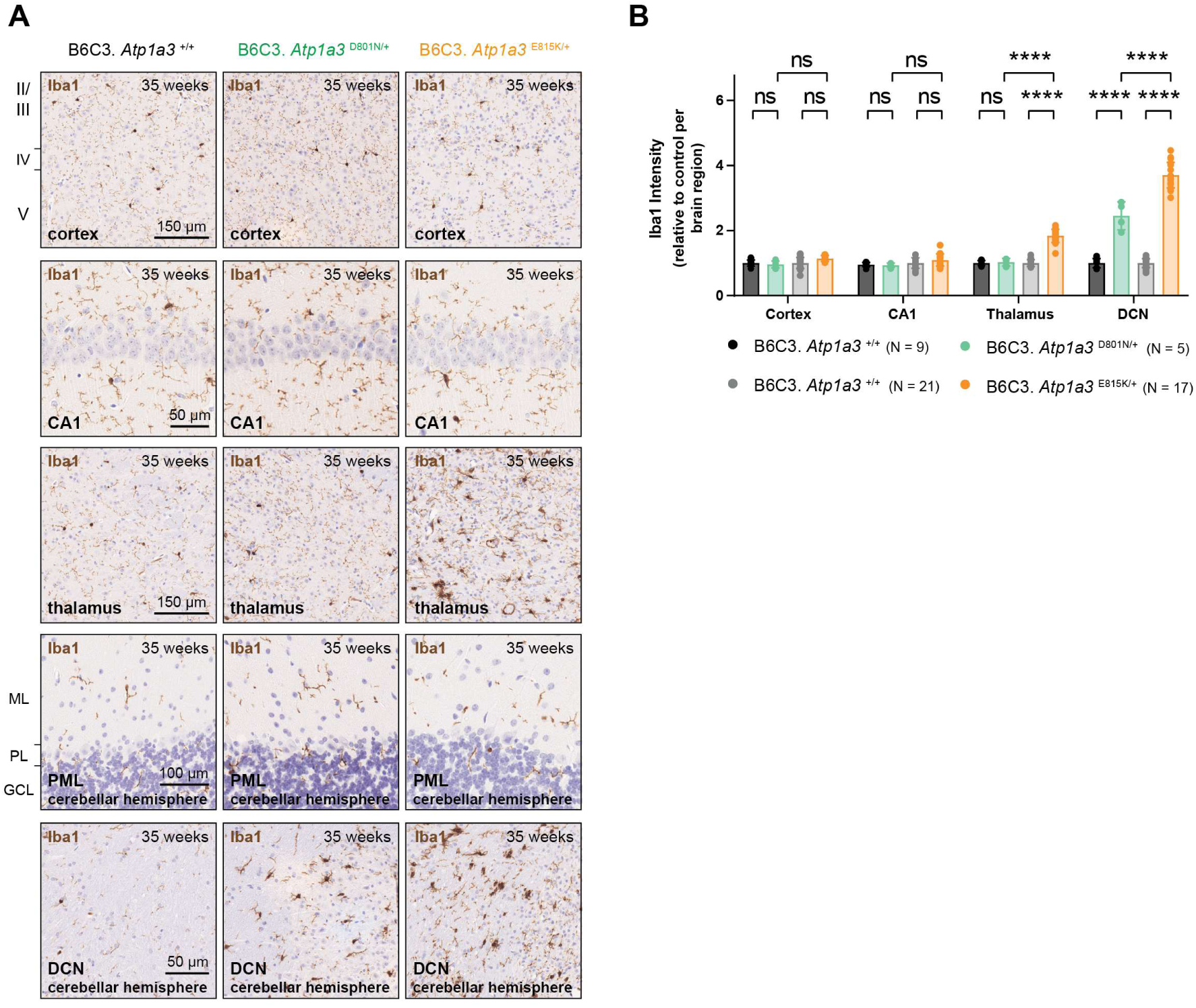
Iba1 analysis in B6C3 AHC mice. (A) Immunohistochemistry using antibodies to ionized calcium-binding adapter molecule 1 (Iba1, microglia, brown) on parasagittal brain sections from 35-week-old wild type, B6C3.*Atp1a3* ^D801N/+^ and B6C3.*Atp1a3* ^E815K/+^ mice. Sections were counterstained with hematoxylin and higher magnifications of individual brain regions are shown. (B) Relative intensity of Iba1 of wild type, B6C3.*Atp1a3* ^D801N/+^ and B6C3.*Atp1a3* ^E815K/+^ mice. Levels are relative to that of controls (wild type) from each cross and brain region. Wild type littermate controls for each mutant strain are shown (black and grey). Data represent mean ± SD. Scale bars: 150 μm (cortex), 50 μm (CA1), 150 μm (thalamus), 100 μm (cerebellar hemisphere, PML) and 50 μm (cerebellar hemisphere, DCN). CA1, hippocampal cornu ammonis area 1; ML, molecular cell layer; PL, Purkinje cell layer; GCL, granule cell layer; PML, paramedian lobe; and DCN, deep cerebellar nuclei. Two-way ANOVA was corrected for multiple comparisons using Tukey method (B). ns, not significant; **** p ≤ 0.0001. Related to Figure 4.

## Materials and Methods

### Mouse strains

Several mouse strains have been generated as part of this study and the strain details are outlined below. For the generation of B6J-*Atp1a3* ^D801N/+^ mice (C57BL/6J-*Atp1a3*^em3Lutzy^/Lutzy or *Atp1a3* ^D801N, L802L^), plasmids encoding a signal guide RNA to introduce the D801N point mutation in exon 17 of the *Atp1a3* gene and the Cas9 nuclease were introduced into the cytoplasm C57BL/6J-derived fertilized eggs (B6J, JR #664, The Jackson Laboratory). An additional silent nucleotide change (CTG to CTA, L802L) was introduced as part of the CRISPR/Cas9 inactivating strategy. Targeted heterozygous embryos were transferred to pseudo-pregnant females. Pups were analyzed for correct targeting and the absence of off-targets by sequencing and PCR for further breeding. Survival data for this strain were previously provided to support other ATP1A3 research studies (Y. B. Liu et al., 2024). The congenic B6.129(Cg)- *Atp1a3^tm1.1Mika^*/Mmjax (also known as ‘Matb’ or E815K *knock in* mice) were previously generated and have been kindly provided by Dr. Mohamad A. Mikati (Helseth et al., 2018) (MMRRC #069591 at The Jackson Laboratory, MGI:6856863).

Because the congenic B6J AHC mice were highly susceptible to early, sudden and/or unexpected death, and hindered the maintenance of either strain, we transferred the corresponding AHC mutations onto a hybrid vigor genetic background using *in vitro* fertilization (IVF). For the generation of B6C3.*Atp1a3* ^D801N/+^ mice (B6C3-*Atp1a3*^em3Lutzy^/Mmjax), an IVF expansion utilizing sperm from heterozygous C57BL/6J- *Atp1a3*^em3Lutzy^/Lutzy males and C3H/HeJ (C3H, JR #659, The Jackson Laboratory, MGI:2159741) females as oocyte donors was performed and produced wild type (B6C3.*Atp1a3* ^+/+^) and heterozygous mutant (B6C3.*Atp1a3* ^D801N/+^) mice. To ensure sufficient heterozygosity between the B6J and C3H genetic backgrounds for strain maintenance, heterozygous B6C3.*Atp1a3* ^D801N/+^ males are crossed to B6C3F1/J (JR #100010, The Jackson Laboratory, MGI:5654213) females every generation and the resulting progeny are subject to experimental testing. This strain has been donated to the MMRRC at the Jackson Laboratory and is publicly available (MMRRC #071287 at The Jackson Laboratory, MGI:7461667).

For the generation of B6C3.*Atp1a3* ^E815K/+^ mice (*Atp1a3*^tm1.1Mika^/LutzyMmjax), an IVF expansion utilizing sperm from heterozygous B6.129(Cg)-*Atp1a3^tm1.1Mika^*/Mmjax (MMRRC #069591 at The Jackson Laboratory, MGI:6856863) males and C3H/HeJ (C3H, JR #659, The Jackson Laboratory, MGI:2159741) females as oocyte donors was performed and produced wild type (B6C3.*Atp1a3* ^+/+^) and heterozygous mutant (B6C3.*Atp1a3* ^E815K/+^) mice. To ensure sufficient heterozygosity between the B6J and C3H genetic backgrounds for strain maintenance, heterozygous B6C3.*Atp1a3* ^E815K/+^ males are crossed to B6C3F1/J (JR #100010, The Jackson Laboratory, MGI:5654213) females every generation and the resulting progeny are subject to experimental testing. This strain has been donated to the MMRRC at the Jackson Laboratory and is publicly available (MMRRC #071376 at The Jackson Laboratory).

Upon discovery of the reduction in fragility of B6C3 AHC mice and to support potential applications for other ATP1A3 researchers, we publicly shared this information and data with the ATP1A3 and AHC community through the annual ATP1A3 symposium in 2022 and 2023, and the public mouse strain catalog of the Jackson Laboratory. Mice were genotyped at birth, wean age (∼postnatal P28), and triple confirmed when they reached the expected study end point (e.g., necropsy for tissues collection and/or histology). The genotyping primers are listed below for each mutant allele and are applicable regardless of the genetic background. The Jackson Laboratory Animal Care and Use Committee, the ethical committee of the Leiden University, in accordance with ARRIVE guidelines and EU Directive approved all mouse protocols.

A) D801N mutation

Common Forward: 5’ CTC TTG GCA CCA TCA CCA TC’3

Common Reverse: 5’ TTT AGT AGC AGC CAG GCT TAC C’3 WT

Probe (HEX): 5’ CTG CAT TGA CCT GGG TAC C’3

D801N mutant Probe (FAM): 5’ TCT GCA TTA ACC TAG GTA CCG AC’3

B) E815K mutation

Common Forward: 5’ ACT CAC ACA GCC TGC CTC TC’3

Common Reverse: 5’ CCT CTT CAT GAT GTC GCT CTC’3 WT

Probe (HEX): 5’ TGG CCT ACG AGG CTG C’3

E815K mutant Probe (FAM): 5’ TGG CCT ACA AGG CTG CC’3

### Mouse cohorts

Several cohorts of mice were generated as part of this study and the details are outlined below. All testing was done during daylight hours and performed using mice of either sex.

1. *Body weight cohort*. Body weights shown were recorded weekly starting from birth up to ∼32 weeks of age. Mice were not subjected to any other behavioral testing.
2. *HIP cohort*. HIP experiments shown were performed on mice at 6 and 12 weeks of age. Mice were not subjected to any other behavioral testing.
3. *Rotarod cohort*. Rotarod experiments shown were longitudinally performed on mice at 10 and 12 weeks of age. Mice were not subjected to any other behavioral testing.
4. *Open field cohort*. Open field experiments shown were performed on mice at 15 weeks of age. Mice were not subjected to any other behavioral testing.
5. *Blood collection, grip strength, rotarod and necropsy cohort*. Retro-orbital blood collection (for NFL analysis) was performed on mice at various ages, grip testing at 16 weeks of age, rotarod testing at 24 weeks of age, and then, were collected for necropsy at 35 weeks of age. Mice were not subjected to any other behavioral testing.
6. *ECG cohort*. Mice were tested at 22 weeks of age for EKG testing.
7. *Electrophysiology cohorts*. Mice were longitudinally subjected to electrophysiological testing (e.g. epileptiform activities, PSD analysis and spontaneous SD) at 6, 10 and 14 weeks of age. Mice were not subjected to any other behavioral testing.
8. *Molecular biology cohorts*. Tissue samples from mice were collected for molecular analysis. Mice were not subjected to any other behavioral testing.
9. *DIVA cohorts*. Mice were longitudinally monitored between 13 to 22 weeks of age. Mice were not subjected to any other behavioral testing.
10. *Survival*. The survival of all generated and sampled mice (birth to pre-defined study end) was monitored. The survival of B6C3 AHC mice has not been monitored and/or tested past 8.5 months of age. The survival data does not include data of breeders because the recording of litters of the breeder colony only occurred between P10-P23, only male mice were sampled and genotyped, and male breeders were replaced at ∼14- 16 weeks of age to refresh breeding units. The survival of mice subjected to electrophysiology studies is not included because some but not all mice were internationally shipped, and test subjects required brain surgery for the probe implantation.

## Behavioral and experimental testing

### Survival

In addition to the routine welfare check, animals were monitored (2-3 times per week) from birth on. For spontaneous deaths, mice were found dead without any prior indication of welfare and/or health concerns. In addition, mice were monitored for their overall health by assessing the body condition score (BCS) (Foltz & Ullman-Cullere, 1999). In the presence of a lowering BCS, mice were provided with a diet gel (Nutri-Gel, Bio-Serv) aiding to support health and the overall body condition. Nevertheless, mice that reached BCS ≤ 2, the defined humane end point, were recorded as ‘death’ and included in the survival curve. Although spontaneous and unexpected deaths were observed in B6C3.*Atp1a3* ^E815K/+^ mice, they required humane euthanasia as the study end point (‘death’) due to low BCS. Thus, survival data include spontaneous death and death due to low BCS.

### Hypothermia induced paroxysmal episodes (HIP)

We adopted the previously described hypothermia paradigm with some modifications (Isaksen et al., 2017; Pizoli et al., 2002). A clear plastic box (30 x 30 x 14 cm with a floor area of 728 cm^2^) was filled with 5°C ± 0.2°C cold water (monitored via thermometer) up to a height of ∼4 cm to ensure that the core of the mouse body is submerged in the water but the mouse retains the ability to touch the floor of the box with at least their hind limbs and therefore, avoid the risk of drowning. The body weight and rectal body temperature were measured. The ‘duration in water’ was determined based on body weight (see table below) to accomplish a comparable body temperature reduction of at least 10°C. Note: The body weight difference between wild type and mutant mice becomes increasingly bigger with age and accommodating a comparable reduction in body temperature between genotypes would have required mice (e.g., wild type) to exceed the ‘duration in water’ past 5 minutes. Thus, HIP experiments were not performed on mice past 12 weeks of age to avoid potential welfare and health concerns.

| Sex | Body weight | Duration in water |
| --- | --- | --- |
| Females/Males | Less than 20g | 2 minutes |
| Females/Males | 20 - 30g | 3 minutes |
| Females/Males | More than 30g | 5 minutes |

While restrained, the mouse was transferred into the water and then slowly released (timer to start ‘duration in water’). Subsequently, the mouse was taken out of the water bath, excess of water removed through a paper towel, the rectal body temperature was immediately measured (‘0 minutes’), and the mouse was placed in the ‘recovery station’ for the observation of paroxysmal episodes. One side (floor area of 333 cm^2^) of a duplex cage functioned as the ‘recovery station’ (one mouse per ‘recovery station’). The ‘recovery station’ was on top of a heating pad, which was set to 37°C. Half of the floor of the ‘recovery station’ was covered with cage bedding while the other half remained bare of bedding. The mouse was placed on the bare side to avoid the cage bedding causing a visual obstruction for the operator (video example, see Supplementary file 3). Upon recovery of mobility, mice generally moved into bedding (e.g. hiding, grooming, ….). The rectal body temperature was measured at 15-, 30- and 60-minutes post water exposure. The mouse was moved from the ‘recovery station’ back into the home cage upon complete recovery (body temperature, normal mobility and mouse behavior). The scoring of paroxysmal episodes (dystonia- and convulsion-like events) was performed post water exposure (start of ‘recovery period’) and we adopted a scoring system (see table below) with some modifications based on previous studies (Pizoli et al., 2002).

| Dystonia-like events |  |  | Convulsion-like events |
| --- | --- | --- | --- |
| HIP Score | Pizoli et al, 2002 | This study | This study |
| 0 | No motor abnormalities and no dystonia | No motor abnormalities and no dystonia | Recorded as "Yes" or "No" independent of dystonia-like events (score) |
| 1 | Slowed motor behavior but no dystonia | Slowed motor behavior but no dystonia |  |
| 2 | Limited movement/ambulation, dystonic posture/event when disturb | Dystonia only occurs when the mouse is "<u>disturbed</u>" such as handling during body temperature measurement (distinction to mere "hypothermic" events) |  |
| 3 | Moderate impairment of movement, frequent and spontaneous dystonic events/postures | Dystonia occurred, animals <u>retain ability</u> to hold/balance/right themselves, dystonic events may occur frequent/multiple times |  |
| 4 | Severe impairment of movement, sustain and frequent dystonic events/postures | Dystonia occurred, animals <u>occasionally retain ability</u> to hold/balance/right themselves, dystonic events are longer lasting and may occur frequent/multiple times |  |
| 5 | Prolonged immobility in dystonic postures/events | Dystonia occurred, animals <u>fail to retain ability</u> to hold/balance/right themselves, e.g. lay flat on stomach, back or side, no signs to correct position, dystonic events are long lasting |  |

Note: Since Pizoli et al, 2002 included the option that defects may occur upon ‘disturbance’ (score 2), we wanted to allow for the possibility that hypothermia on its own is not sufficient to induce events but instead also requires further ‘disturbance’. Therefore, we defined the score of 2 as dystonia-like events that occur upon ‘disturbance’. Because operators are required to record the body temperature (physical handling) at consistent and pre-defined time intervals for any mouse, we utilized this as ‘disturbance’. Thus, a score of 2 would only be recorded if events immediately followed the measurement of body temperature. However, we did not observe such events and therefore, our data lack observations for a score of 2 (Figure 1D).

### Rotarod

Mice were habituated to the testing room one hour prior to testing with the rotarod on and rotating (Ugo Basile Rotarod for Mouse). The apparatus was cleaned before and in between each mouse with 70% ethanol. Mice were tested for four consecutive trials of accelerating rotarod starting at 5 RPM and ramping up to 40 RPM over 300 seconds. Latency to fall for each of the four trials was recorded with a 45 second rest period between each trial. Day one is considered training day, followed by the same protocol on the subsequent day which is considered testing day (Figure 2A). The latency from the testing day trials 2, 3, and 4 were averaged and reported as latency to fall for each mouse. For the assessment of motor learning function (Figure S2A), the exact same protocol was used with additional, consecutive test days (1 training day and 3 testing days).

### Grip strength

A commercially available grip-strength meter (Bioseb) attached to a wire grid was used to measure front paw and hind paw peak grip strength (kg). Mice were weighed and allowed to acclimate to the testing room for a minimum of 60 minutes before testing. For fore paw testing, mice were held by their tails and lowered toward the grid to allow for visual placing and for the mouse to grip the grid with its forepaws. The animal was firmly pulled horizontally away from the grid. With hind paw testing, each mouse was restrained, and hind paws were dragged the length of the grid and peak grip strength for was recorded. An average of 3 trials served as the final peak force. The grip strength is shown with and without normalization to the mouse body weight.

### Open field

Open field arenas used in this study were squares (40 x 40 x 40 cm) and made from clear Plexiglass. LED lights indirectly illuminated the arenas at ranges of 100-500 ± 20 lux. The chambers were vented, and sound attenuated. Mice were habituated to the room for 60 minutes prior to the test. One mouse was placed in the center of each arena, and vertical and horizontal activity were recorded by beam breaks across two levels in infrared beams using the Fusion software (OmniTech Electronics). Mice were recorded for 60 minutes, and the arenas were cleaned with 70% ethanol between subjects within a single testing day. Several readouts were obtained from the open field analysis and the details are listed below.

1. *Total distance traveled.* The total distance that the subject has traveled.
2. *Total movement time.* The length of time that the subject spent in activity. Activity is defined as a period in which ambulation and/or stereotypy occurred.
3. *Total rest time.* The length of time that the subject spent at rest. A resting period is defined as a period of inactivity greater than or equal to 1 second.
4. *Stereotypic episode count.* The number of beam breaks due to stereotypic activity. If the animal breaks the same beam (or set of beams) repeatedly then the monitor considers that the animal is exhibiting stereotypy. This typically happens during grooming or head bobbing.
5. *Vertical episode count.* This is incremented by 1 each time the animal rears. The animal must go below the level of the vertical sensor for at least 1 second before the next rearing can be registered.
6. *Total margin time.* Time spent by the animal in proximity to the walls of the cage. This area is defined as the exterior 4x16 and 16x4 matrices for the left right bottom and top regions.
7. *Total center time.* The total time spent in the center portion of the cage. This area is defined as the center 8x8 square matrix plus the coordinates between the outermost beams of the area and the adjacent non-area beams.

### Electrocardiogram

ECG studies were performed as previously described (Oestereicher et al., 2023). Briefly, mice were anesthetized in the induction chamber with 2-3% isoflurane. Once the mouse was in lateral recumbency and unable to right itself, eye lubricant was applied, and the animal was transferred to a nosecone for maintenance with 1-2% isoflurane. Anesthetized mice were positioned supine on a warming pad apparatus that maintained the animal’s core temperature at 37 °C. Surface ECG recording was using leading II configuration. The needle electrodes were placed subcutaneously; the negative electrode in the right forelimb; the ground electrode in the right hindlimb; and the positive electrode in the left hindlimb. ECG was recorded in a dimly lit, quiet procedure room. A steady heart rate within the typical mouse range was monitored to ensure that the appropriate anesthetic depth was reached. In order to eliminate circadian influences, ECG was recorded during the morning when the resting phase of a mouse begins. ECG data were collected for up to 120 s and the resulting data analyzed using LabChart software (ADInstruments). The P, Q, R, S and T markers were identified and adjusted as follows: P Start was positioned immediately before the P-wave dip, P Peak was identified as the highest point of the P wave, and P End was marked where the P wave returned to baseline. QRS Start was placed immediately before the QRS wave dip, QRS Max was designated as the highest point of the QRS wave, and QRS End was located between the lowest amplitude of the QRS wave and the J-wave. ST Height was fixed 10 msec from QRS Max. T Peak was determined as the highest or lowest point of the T wave, depending on its polarity, and T End was marked where the T wave returned to baseline, falling between T Peak and P Start. If T End was not visible, the Maximum RT was adjusted in the settings. The QT intervals were also corrected as QTc using the following “Bazett” method: QTc = QT/(RR/100)^0.5^.

### Surgery for electrophysiology

For long-duration electrophysiological local field potential (LFP) recordings of brain activity, 5- to 6-week-old (male and female) B6C3.*Atp1a3* ^D801N/+^ (D801N) (N = 7), and B6C3.*Atp1a3* ^E815K/+^ (E815K) (N = 11) and B6C3.*Atp1a3* ^+/+^ (wild type) (N = 5 for each strain; data were combined in the analyses) mice underwent surgery under isoflurane anesthesia (induction 4%; maintenance 1.5%) with oxygen air enrichment and body temperature maintained at 37 ± 0.5°C. LFP electrodes (75 μm platinum(Pl)/iridium(Ir); PT6718; Advent Research Materials, Oxford, UK) were implanted in the following configuration (in mm relative to bregma, anterior/lateral/ventral, respectively; see also schematic in Figure 3A): bilateral visual cortex (V1: -3.5/2.0/-0.5), bilateral dorsal hippocampus (dHC: -2.0/1.5/-1.3) and unilateral (right) motor cortex (M1: +1.5/1.8/-0.5). For all operated mice, two low-impedance 75 µm Pt/Ir electrodes with ∼1 mm uninsulated tip were also placed in the cerebellum, as reference and ground. All electrode locations were post hoc histologically verified. The recording electrodes were connected to a 7-channel pedestal (E363/0 socket contacts and MS373 pedestal; Plastics One, Roanoke, VA, USA) and connected to the skull using dental cement (075794; Sun Medical SuperBond C&B with L-polymer; Hofmeester, The Netherlands). Carprofen (5 mg/kg, s.c.) was given for post-operative pain relief. All mice survived the surgical procedure and showed no discomfort from surgery during recovery.

### Long-duration local field potential recordings

Following a post-surgery recovery of at least 24 hours, the mice were transferred to shielded EEG recording cages in which animals could behave freely. Mice were connected to the EEG system at 6, 10 and 14 weeks of age and recorded for 3-5 days at each given age. The system consisted of a Faraday cage, in which the mouse was connected to custom-built recording hardware via a custom-made counterbalanced, low-torque commutator. LFP recordings were 3X pre-amplified, filtered and fed into separate amplifiers for direct current (DC)-potential (10X gain, relative to reference; DC-500 Hz) and alternating current (AC)-potential signals (200X gain for hippocampal and 400X gain for cortical recordings, relative to reference; 0.05- 500 Hz). Recordings were digitized (Power 1401 and Spike2 software, CED, Cambridge, UK) at a sampling rate of 1000 Hz for the DC-potential and at 5000 Hz for AC-potential signals.

### Electrophysiological data analysis

Electrophysiological AC-LFP recordings and DC-potentials were analyzed off-line using Spike2 software (CED, Cambridge, UK), and custom-written Python and MATLAB scripts. To assess spiking activity including spike-wave discharges (SWDs), isolated hippocampal spikes, and giant spikes, AC-potential signals of LFP recordings from cortex (V1 and M1) and hippocampus in a 24-hour window were analyzed. The window started at least 24 hours after the start of the recording to avoid novelty or stress-induced changes related to connecting the mouse to the EEG recording system. AC-potential LFP recordings were artifact-rejected, band-pass-filtered (0.5 - 100 Hz) and down-sampled to 1000 Hz. Spike-wave discharges (SWDs) were quantified from the frontal M1 LFP channels based on previous criteria (Kros et al., 2015; Letts et al., 2014). In short, after automatic spike detection, a SWD was defined as an asymmetric complex of at least 3 cycles of sharp negative-positive going spikes and waves with peak-to-peak amplitudes at least two-fold higher than background, a minimal discharge frequency of 6 Hz, a minimal duration of 1 second and inter-SWD episode interval ≥ 1 second. To detect hippocampal isolated spikes and giant spikes, first, an automatic spike detection method was employed using the Nonlinear Energy Operator (NEO) combined with Automatic NEO Thresholding (Yang & Mason, 2017), after which hippocampal giant spikes were identified and extracted from the total number of spikes. Giant spikes were defined as simultaneous spike (either positive or negative) present in all channels, with a large amplitude (> ±10 SD from the filtered baseline) in at least one hippocampal channel, followed by positive deflection (‘after hyperpolarization’) lasting > 200 ms across all channels (Gureviciene et al., 2019). Next, remaining detected hippocampal spikes were classified as ‘isolated hippocampal spikes’ based on their features fitting the criterion of a negative-going brief (< 100 ms) single spike, without a corresponding spike within a ±100-ms window in either the contralateral hippocampus or cortex. The frequency of SWDs, isolated hippocampal spikes and giant spikes was reported as the number of events per hour across a 24-hour period of LFP recording, without correlating the data to specific vigilance states.

To assess occurrence of spontaneous cortical and hippocampal spreading depolarization (SD) events, the entire period of available LFP recordings was analyzed. Cortical or hippocampal SD events were defined in DC-potential recordings as transient negative DC shifts with an amplitude > 5 mV, detected with a subsequent delay between at least two LFP recording electrodes associated with a suppression in AC amplitude as visible in the AC-LFP sonogram. Burst spiking activity preceding SD events was defined as high-amplitude (> 2X value of averaged baseline root mean square of the AC-LFP signal) rhythmic discharges that clearly represented an abnormal EEG pattern (typically consisting of repetitive high-amplitude spikes and slow waves), lasting for ≥ 5 seconds, followed by suppression of AC-LFP activity. The duration of burst activity was determined from the dorsal HC burst by measuring the interval between the first and last spike of a burst. Intra-burst spiking frequency was calculated by dividing the total number of spikes by the duration of the burst. SD amplitude was quantified through identifying the lowest of the depolarization during the SD shift, while the 50% SD-recovery time was assessed from the start of the SD (i.e., start of DC-deflection) to the time-point at which the SD shift had recovered to half of its amplitude level.

To assess cortical network activity features including power spectral analysis (PSD) during active and quiet wakefulness, LFP recorded during the morning (6-12 AM), without the presence of SD events, was analyzed. Vigilance state was determined per 5-second epoch using V1 cortical LFP, and the reference signal and locomotor activity were recorded by the infrared motion detection (PIR) sensor. Active wakefulness was defined by a θ-δ power ratio of > 2 during epochs containing locomotor activity and/or high variance in the reference signal. Quiet wakefulness was identified by the presence of desynchronized LFP with no locomotor activity and a stable reference signal, in between epochs of active wakefulness. To differentiate quiet wakefulness from in particular non-rapid-eye-movement (NREM) epochs, the latter were determined by the presence of high-amplitude δ activity indicative of slow-wave-sleep. When quiet wakefulness lasted > 20 seconds and transitioned into NREM sleep, the first 20 seconds were defined as transition states (i.e., wakefulness to sleep) and not included in the analysis. Attention was given to excluding any large artefact-related activity that could influence the LFP recordings, ensuring accurate assessment of vigilance states. When there was doubt about a sleep - awake period, video footage was used for validation. Next, epochs identified for active and quiet wakefulness were screened for noise deflection and spike exclusion. In total, 5 minutes of spike-free randomly selected epochs during active and quiet wakefulness were used for θ peak frequency and PSD analysis. Power spectra were computed from V1 cortical LFP by applying a Hamming window over each 5-second epoch, followed by a fast Fourier transform and averaging of the resulting power spectra. For the power comparison of specific frequency bands, oscillations were defined and averaged across the mouse-specific bands: delta (1-5 Hz), theta (5-10 Hz), alpha (10-13 Hz), beta (13-30 Hz) and gamma (30-100 Hz).

### Tissue collection

Animals were euthanized by CO_2_ asphyxiation and tissues were harvested, frozen on dry-ice, and stored in -80°C freezer until use.

### Detection of Na^+^/K^+^ ATPase activity and western blotting

Hippocampal tissues from naïve 10-week-old B6C3.*Atp1a3* ^+/+^ (wild type), B6C3.*Atp1a3* ^D801N/+^ (D801N), and B6C3.*Atp1a3* ^E815K/+^ mice (E815K) were homogenized in a buffer of 315 mM sucrose, 20 mM Tris, 1 mM EDTA, pH 7.4, supplemented with complete mini protease inhibitor cocktail (Roche Diagnostics GmBH, Germany). Protein concentrations in crude membrane homogenates were determined with the Lowry method. Gel electrophoresis was performed using mPage 4-12% MES SDS gels (Millipore, USA) followed by transfer onto nitrocellulose membranes (Bio-Rad, USA). Blots were probed with the mouse monoclonal anti-ATP1A3 (α3) antibody F-1 (Santa Cruz Biotechnology, USA). Anti-GAPDH-HRP antibody (Cell Signaling Technology, USA) was used as loading control. Signals were developed with WesternBright ECL reagents (Advansta, USA) and quantified with a GE Healthcare LAS 4000 imaging system and ImageQuant software.

ATPase activity was determined by quantifying hydrolysis of ATP. In order to open vesicular structures formed during homogenization, crude membrane homogenates were pre-treated with SDS (final detergent: sample protein ratio of 0.58). The Na^+^/K^+^-ATPase is resistant to denaturing effects of SDS at this ratio. BSA was added to the solution in order to buffer the exposure to detergent (Kathleen J Sweadner, 2016). Briefly, hippocampi homogenates were preincubated with SDS for 10 minutes at room temperature in a solution containing 2.0 mg/ml BSA. The reaction was stopped by addition of three volumes of a solution containing 0.3 mg/ml BSA and no additional detergent. The final ATPase reaction mixture contained 140 mM NaCl, 20 mM KCl, 4 mM MgCl2, 3 mM disodium ATP (Sigma, vanadate free), 30 mM histidine, pH 7.2, in the presence of different concentrations of ouabain, the specific inhibitor of Na^+^/K^+^-ATPase. The reaction was incubated for 15 minutes at 37 °C, followed by quenching with acid-molybdate and addition of Fiske-Subbarow reducing reagent. Developed color was read at 700 nm. The reaction was performed either in duplicate or triplicate. Na^+^/K^+^-ATPase due to ATP1A3 (α3) was defined as the difference between total activity with no inhibitor added and the one inhibited by 10 μM ouabain, whereas activity due to ATP1A1 (α1) as that active in 10 μM ouabain but inhibited by 3 mM ouabain. The data are expressed as µmol of inorganic phosphate released by 1 mg of total protein in an hour (µmol/mg/hour). There are three isoforms of catalytic subunit of Na^+^/K^+^-ATPase in brain. α1 is expressed widely, whereas ATP1A2 (α2) and ATP1A3 (α3) are specific for glia and neurons, respectively. Activity conferred by ATP1A2 (α2) is no more than 5-10% of the total activity and it may be ignored (Y. B. Liu et al., 2024). In mice, α1 and α2+α3 isoforms differ in their sensitivity to ouabain and this allows to separation of effects on ATPase activity conferred by α1 and α2+α3 isoforms (Figure 1C). Most of the Na^+^/K^+^-ATPase activity recovered from hippocampal samples came from the α3 isoform. Quantifications were performed using mice of either sex.

### Reverse transcription and quantitative PCR (TaqMan) analysis

Mouse tissues (cortex, hippocampus, brainstem and cerebellum) were collected from naïve 10-week-old B6C3.*Atp1a3* ^+/+^ (wild type), B6C3.*Atp1a3* ^D801N/+^ (D801N), and B6C3.*Atp1a3* ^E815K/+^ mice (E815K) with four biological replicates per genotype and brain region. The RNA was isolated using RNeasy Mini Kit and DNase-treated following the manufacturer’s protocol (Qiagen) and was used for cDNA synthesis using High-Capacity cDNA Reverse Transcription Kit and oligo dT primer (ThermoFisher) as per the manufacturer’s protocol. Quantitative RT-PCR (TaqMan) reactions were performed using the Taqman Fast Advance MasterMix (ThermoFisher) and the QuantStudio Flex 7 qPCR detection system (ThermoFisher). For quantitative expression analysis, TaqMan probes (see table below) were multiplexed with probes for the gene of interest and input control (*Tata-binding protein, Tbp*). The expression levels of the gene of interest (FAM) were normalized to those of *Tbp* (HEX) using the 2-ΔΔCT method (Livak & Schmittgen, (A) 2001) and expressed as the fold change ± standard error of the mean (SEM) relative to wild type. Quantifications were performed using mice of either sex.

TaqMan Probes:

| Gene name | Species | Exon location | Fluorophore | Vendor | Cat. number |
| --- | --- | --- | --- | --- | --- |
| *Bmp3* | Mouse | 2 - 3 | FAM | IDT | Mm.PT.58.11601208 |
| *Olfm4* | Mouse | 2 - 3 | FAM | IDT | Mm.PT.58.14228836 |
| *Htra4* | Mouse | 3 - 4 | FAM | IDT | Mm.PT.56a.6274186 |
| *Rara* | Mouse | 6 - 7 | FAM | IDT | Mm.PT.58.11675696 |
| *Crhr2* | Mouse | 6 - 7 | FAM | IDT | Mm.PT.58.12499462 |
| *Th* | Mouse | 12 - 13 | FAM | IDT | Mm.PT.58.33106186 |
| *Itgax* | Mouse | 5 - 6 | FAM | IDT | Mm.PT.58.42516719 |
| *Car9* | Mouse | 1 - 3 | FAM | IDT | Mm.PT.58.14228651 |
| *Tbp* | Mouse | 4 - 5 | HEX | IDT | Mm.PT.39a.22214839 |

### RNA sequencing library construction

Mouse tissues (cortex, hippocampus, brainstem and cerebellum) were collected from naïve 10-week-old B6C3.*Atp1a3* ^+/+^ (wild type), B6C3.*Atp1a3* ^D801N/+^ (D801N), and B6C3.*Atp1a3* ^E815K/+^ mice (E815K) to obtain four (2 female and 2 male mice) biological replicates per genotype and brain region. The RNA was isolated using RNeasy Mini Kit and DNase-treated following the manufacturer’s protocol (Qiagen) and used for RNA library construction as per the manufacturer’s protocol (KAPA mRNA HyperPrep Kit, Roche Sequencing and Life Sciences). Briefly, the protocol includes isolation of polyA containing mRNA using oligo-dT magnetic beads, RNA fragmentation, first and second strand cDNA synthesis, ligation of Illumina-specific adapters containing a unique barcode sequence for each library, and PCR amplification. The quality and concentration of the libraries were assessed using the D5000 ScreenTape (Agilent Technologies) and Qubit dsDNA HS Assay (ThermoFisher), respectively. All libraries were pooled (48 samples), and the pool was sequenced as paired end (150bp) on an Illumina NovaSeq X Plus using the 10B Reagent Kit with a target of at least 30 million reads per sample.

### Data analysis of RNA sequencing data

First, adapters were trimmed from read pairs using cutadapt (Martin, 2011) version 4.4 using parameters - a AGATCGGAAGAGCACACGTCTGAACTCCAGTCA-A AGATCGGAAGAGCGTCGTGTAGGGAAAGAGTGT and -m 40. Trimmed reads were used to quantify expression with kallisto version 0.48.0 (Bray et al., 2016) by pseudo-aligning to a GENCODE version M35 reference transcriptome (with --bias parameter set). The resulting transcript abundance estimates were imported into R using tximport (Soneson et al., 2015) for differential expression analysis using DESeq2 version 1.38.3 (Love et al., 2014). For each tissue, expression of B6C3.*Atp1a3* ^D801N/+^ (D801N) or B6C3.*Atp1a3* ^E815K/+^ mice was compared to expression of B6C3.*Atp1a3* ^+/+^ (wild type) using a linear model including sex as a covariate and testing for differences in genotype. Fold change was estimated using an adaptive shrinkage estimator from the ahsr (version 2.2-65) R package (Stephens, 2017).

Codes for the data analysis, quality control, and data processing have been made available at https://github.com/scottiadamson/atp1a3_mouse_rna_seq_analysis

### Pathway analysis

Data were analyzed using Ingenuity Pathway Analysis (IPA, QIAGEN Inc., https://www.qiagenbioinformatics.com/products/ingenuity-pathway-analysis). Kyoto Encyclopedia of Genes and Genomes (KEGG) pathway analysis was performed using the ShinyGO v0.61 bioinformatics web server (http://bioinformatics.sdstate.edu/go) (Ge et al., 2020) by uploading the gene lists from our RNA sequencing analysis. KEGG pathway terms with a *p*-value cutoff (FDR) ≤ 0.05 were considered enriched.

### Histology and immunohistochemistry

Mice were transcardially perfused with PBS and then with 10% Neutral buffered formalin (NBF) for immunohistochemistry (IHC). Brain tissues were post-fixed (24 - 48 hours) in NBF and then embedded in paraffin. For IHC staining, sections were deparaffinized, rehydrated, and the NBF-fixed, paraffin embedded sections were stained with antibodies against GFAP (Abcam 16997, 1:100, antigen retrieval via citrate buffer for 20 minutes) and Iba1 (Abcam 178846, 1:2000, antigen retrieval via citrate buffer for 5 minutes) on the Leica-Bond auto-staining system (Leica Biosystems). Sections were counterstained with Hematoxylin and histological slides were digitalized at 40x resolution using a digital slide scanner (Hamamatsu NanoZoomer). For GFAP and Iba1 quantification, the staining intensity was measured in an area of 0.3 mm^2^ (somatosensory cortex, layer I to VI), 0.09 mm^2^ (CA1 of hippocampus), 0.5 mm^2^ (thalamus), and 0.1 mm^2^ (DCN of the cerebellum) using ImageJ, averaged from three parasagittal brain sections (three brain sections spaced 150 μm apart) per animal and expressed as the fold change relative to wild type mice per strain and brain region. Quantifications were performed using mice of either sex.

### Serum collection and NFL analysis

Whole blood was collected via retro-orbital collection (RO) by applying a topical anesthetic (Proparacaine) on the eye. The collected blood was transferred into the serum collection tube (Microtainer Tube with serum separator, yellow cap), the tube was inverted a few times, kept at room temperature for 20-30 minutes, stored on wet-ice and then, centrifuged for 10 minutes at 14,000 x g (rcf) at 4°C. Afterwards, the serum located above the gel separator matrix was transferred into a new storage tube and stored at -80°C until further analysis. Serum samples were subjected to neurofilament light chain (NFL) analysis with a 32x sample dilution and 2 replicates (technical replicates) per sample via Simoa HD-X analyzer (Simoa NF-Light v2 Advantage Kit #104073 by Quanterix). Quantifications were performed using mice of either sex.

### Statistics

For quantification of RNA expression (TaqMan), ATPase activity and protein expression, mouse behavioral data, serum NFL levels, and histological quantifications, p-values were computed in GraphPad Prism or RStudio using either student’s t-test, Mantel-Cox test, Fisher’s exact test, multiple t-tests, one-way ANOVA, or two-way ANOVA and statistical tests were corrected for multiple comparisons as indicated in the figure legends. Quantifications were performed with mice of either sex. The number of biological replicates is depicted in each figure throughout the manuscript. For behavioral (e.g., HIP, rotarod, open field, grip strength and ECG) and molecular (e.g., NFL, TaqMan, IHC) analysis, operators were blinded during testing, data collection and analysis. For the analysis of RNA sequencing and electrophysiology data, operators were unblinded as the identification of changes and abnormalities required knowledge of the respective genotype. Except for the survival curves, data are shown as the pool of ‘male + female’ mice because the absence or presence of significant changes were independent of sex.

## Data availability

The RNA sequencing data have been deposited to GEO with the identifier (GSE279074). Details about the sample and sequencing inventory have been provided (see Supplementary File 6 - Sample Sequencing log).

## Supplementary files

**Supplementary File 1 - DIVA cage B6C3 D801N 14 wks.** Example of home cage behavior of B6C3.*Atp1a3* ^D801N/+^ (D801N) mice. Four heterozygous D801N mice are shown at 14 weeks of age with one mutant mouse developing a lethal paroxysmal episode.

**Supplementary File 2 - DIVA cage B6C3 E815K 19 wks.** Example of home cage behavior of B6C3.*Atp1a3* ^E815K/+^ (E815K) mice. One heterozygous E815K mouse is shown at 19 weeks of age developing a lethal paroxysmal episode.

**Supplementary File 3 - Hypothermia induced paroxysmal spells.** Example of paroxysmal episode and convulsive-like events induced via hypothermia in 12-week-old B6C3.*Atp1a3* ^D801N/+^ (D801N) mice.

**Supplementary File 4 - B6C3 E815K 14 wks.** Example of trembling and unsteady gait movements of 14- week-old B6C3.*Atp1a3* ^E815K/+^ (E815K) mice.

**Supplementary File 5 - Differential gene expression analysis.** Differential expression (DE) of genes from B6C3.*Atp1a3* ^D801N/+^ (D801N) and B6C3.*Atp1a3* ^E815K/+^ (E815K) mice was obtained from different regions of the brain including the BS (brainstem), CM (cerebellum), CX (cortex) and HP (hippocampus). Data were analyzed relative to wild type mice. Positive fold changes refer to an increase in gene expression in mutant mice relative to wild type mice. Negative fold changes refer to a decrease in gene expression in mutant relative to wild type mice. The datasets include the raw *p*-values and the adjusted *p*-values (adj. *p*-value) corrected for multiple comparison.

**Supplementary File 6 - Sample Sequencing log.** Sample and Sequencing inventory of samples subjected to RNA-Sequencing. The log includes details about the samples (e.g., genotype, sex, brain region) and library (e.g., library name, index primer).

**Supplementary File 7 - Neuroinflammatory gene list.** For the comparison of differentially expressed genes in B6C3 AHC mice, we compared data with previously identified neuroinflammatory genes (astrocytes and microglial genes) in mice. Previous studies (Ishimura et al., 2016) consolidated the neuroinflammatory genes derived from multiple studies in mice (Chiu et al., 2013; Holtman et al., 2015; Orre et al., 2014).

## Acknowledgements

We thank the patients and family members of the AHC patient advocacy groups including Nina Frost (Hope for Annabel), Mary Saladino (For Henry), Simon Frost (Cure AHC), Vicky Platt and Stephen Henderson (AHC foundation) for their support and providing the patient perspectives. We thank Dr. Mohamad A. Mikati for sharing the ‘Matb’ (Atp1a3 tm1.1Mika) mice. We thank Drs. Kathryn J. Swoboda, Hendrick Rosewich, Matthew Campbell, Alfred L. George, Christine Q. Simmons, Steven J. Clapcote, Nutan Sharma, Laurie Ozelius, Alexander Souse, and Holt Saki for their clinical and ATP1A3-related research expertise, and valuable discussions. We thank the Scientific Services at the Jackson Laboratory (JAX) including the Mouse Model Generation Core, Reproductive Sciences, Genome Technologies, Center for Biometric Analysis, Clinical Chemistry, Histology and Microscopy Cores for their support with the *in vitro* fertilization, RNA-sequencing, behavioral test equipment, serum NFL analysis, histology and microscopy equipment. We thank the DIVA (Digital In Vivo Alliance) consortium for the access to the digital cage system. We thank all RDTC (Rare Disease Translational Center) and JCPG (JAX Center for Precision Genetics) members for their operational support. This work was supported by the Dutch National Epilepsy Foundation (2022-10; EAT and AMJMvdM), the European Union Joint Programme Neurodegenerative Disease Research project (REBALANCE, 10510062210003; EAT and AMJMvdM), the Medical Delta program “Medical NeuroDelta: Ambulant Neuromonitoring for Prevention and Treatment of Brain Disease” (AMJMvdM), and NIH U54 OD030187 (SAM and CML).

## Author Contributions

MT and LA designed mouse and behavioral experiments. MT and NSM designed the HIP protocol. SP, JS, SM and HJ performed the mouse behavioral experiments with MT’s guidance. PAP, VS, DC, and AD performed molecular experiments of mice with MT’s guidance. CML provided guidance for MT. EA and KJS designed and performed biochemistry experiments. GK, EAT and AMJMvdM designed electrophysiology experiments. GK performed electrophysiology experiments and analyzed data with EAT’s and AMJMvdM’s guidance. MT and SIA designed transcriptome experiments. SIA performed the computational analysis of RNA-sequencing data with DAK’s guidance. SIA guided MT with subsequent data analysis. AZ designed and performed the CRISPR generation of B6J-*Atp1a3* ^D801N/+^ mice. NM, KJS, EAT, AMJMvdM, SAM and CML provided advice for the experiments and the manuscript review. MT wrote the manuscript together with GK and AMJMvdM for the electrophysiology experiments.

